# Balanced release of neuropeptide FF and gonadotropin-releasing hormone 3 modulates male sexual behavior

**DOI:** 10.1101/2021.07.14.452327

**Authors:** Chie Umatani, Nagisa Yoshida, Eri Yamamoto, Yasuhisa Akazome, Yasutaka Mori, Shinji Kanda, Kataaki Okubo, Yoshitaka Oka

## Abstract

Animals properly perform sexual behaviors by using multiple sensory cues. However, neural mechanisms integrating multiple sensory cues and regulating motivation for sexual behaviors remain unclear. Here, we focused on peptidergic neurons, terminal nerve gonadotropin-releasing hormone (TN-GnRH) neurons, which receive inputs from various sensory systems and co-express neuropeptide FF (NPFF) in addition to GnRH. Our behavioral analyses using knockout medaka of GnRH (*gnrh3*) and/or NPFF (*npff*) demonstrated that some sexual behavioral repertories were ‘delayed’, not disrupted, in *gnrh3*^-/-^ and *npff^-/^*^-^ males, while the double knockout showed normal behaviors. We also found anatomical evidence to show that both neuropeptides modulate the sexual behavior-controlling brain areas. Furthermore, we demonstrated that NPFF activates neurons in the preoptic area via indirect pathway, which is considered to induce the increase in the motivation for male sexual behaviors. Considering these results, we propose a novel mechanism by which balanced release of co-existing peptides is important for the neuromodulatory function of TN-GnRH neurons in the control of behavioral motivation. Our results may go a long way toward understanding the functional significance of peptidergic neuromodulation in response to external environments.

## Introduction

When animals mate, they recognize their partners of the opposite sex by some sensory cues, such as chemical and visual ones, and perform sexual behaviors towards them. After their partners of the opposite sex accept their approach, they clasp (or copulate) and reproduce. Although lots of behavioral studies have demonstrated this sequence of sexual behaviors, the neural mechanisms integrating multiple sensory cues and regulating motivation for sexual behaviors remain unclear. In recent three decades, various peptidergic neurons that release neuropeptides have been reported as ones involved in sexual behaviors (Argiolas and Melis, 2013). Among them, we focused on the terminal nerve gonadotropin-releasing hormone (TN-GnRH) neuron for the following reasons. TN-GnRH neurons are suggested to modulate the motivation for sexual behavior, because ablation of the terminal nerve induced decrease in the male sexual behavior (Wirsig and Leonard, 1987, Yamamoto et al., 1997). In addition, this neuron is anatomically suggested to receive sensory information (Yamamoto and Ito, 2000).

Interestingly, TN-GnRH neurons have been found to express neuropeptide FF (NPFF) as a co-transmitter in addition to gonadotropin-releasing hormone (GnRH) (Figure 1A, Oehlmann et al., 2002, Wirsig-Wiechmann and Oka, 2002, Saito et al., 2010; FMRFamide-like peptides are now identified as NPFF). GnRH in the terminal nerve has been suggested to modulate excitabilities (neuromodulation) in the sensory processing neurons (Kinoshita et al., 2007, Kawai et al., 2010, Umatani et al., 2015). In addition, GnRH-induced neuromodulation is suggested to modulate the motivation for sexual behaviors (Li et al., 2017). In rodents, GnRH administration induced both facilitatory and inhibitory effects on sexual behaviors, and the functions of GnRH in sexual behaviors in the vertebrate are somewhat controversial (Dorsa and Smith, 1980, Myers and Baum, 1980, Dorsa et al., 1981, Dennison et al., 1996). On the other hand, the functions of NPFF have only been reported in the spinal cord (Vilim et al., 1999, Yang et al., 2008), even though NPFF and NPFF receptors are expressed in the brain (Gouarderes et al., 2004). Hence, it is still unknown which neuropeptide, GnRH or NPFF in the terminal nerve, plays a key role in sexual behavior.

**Figure 1:**
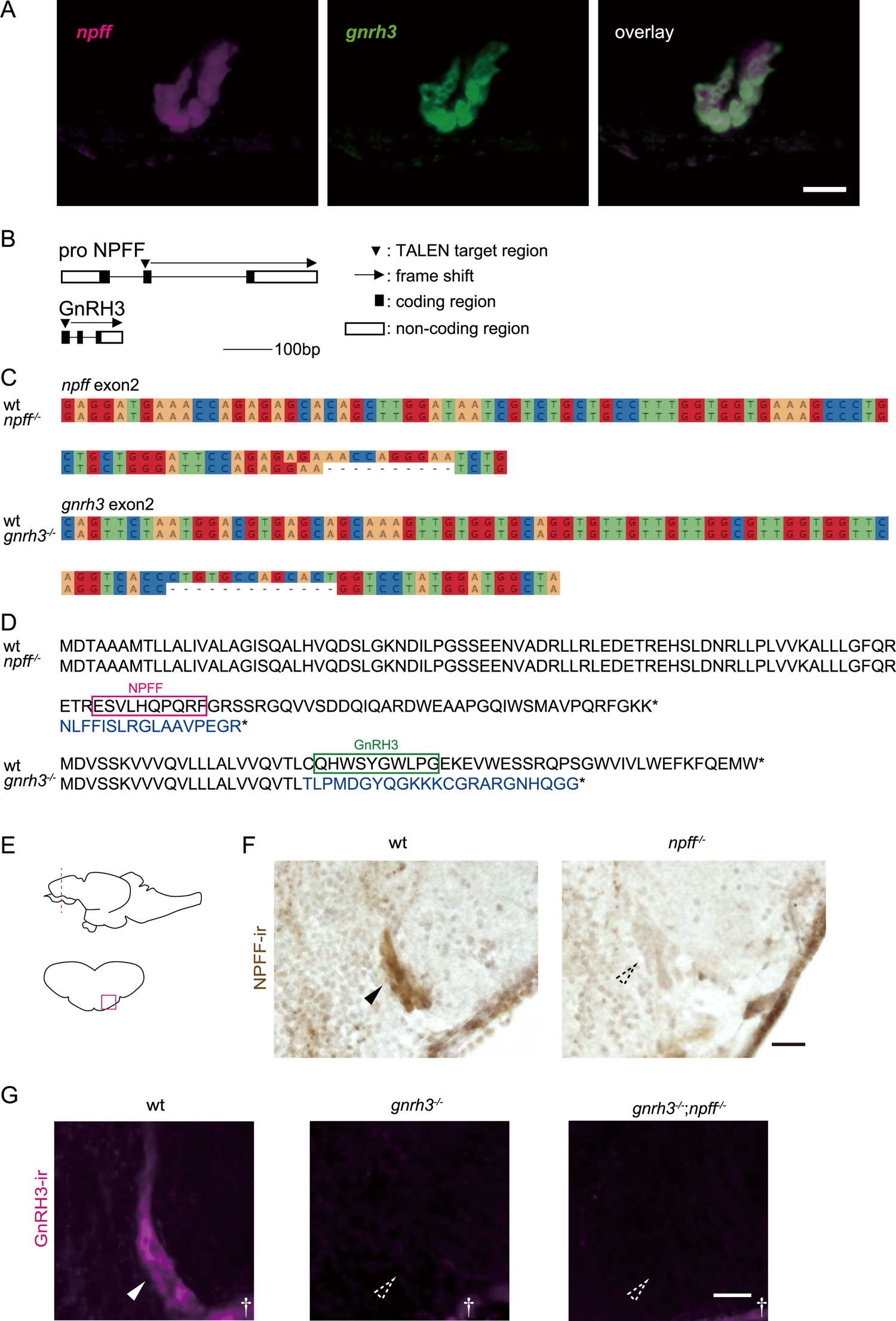
Generation of *npff ^-/-^*, *gnrh3 ^-/-^*, and *gnrh3 ^-/-^*;*npff ^-/-^* medaka (A) *npff* and *gnrh3* are co-expressed in the terminal nerve as demonstrated by ISH. Magenta shows signal for *npff* mRNA, and green shows signal for *gnrh3* mRNA. Scale bar: 30 μm. (B) Schematic diagram of TALEN target site. Black box: coding region, white box: non-coding region of exon, arrowhead: TALEN target region, arrow and dotted line: region where frame shift occurred. (C) The sequence of *npff* or *gnrh3* of knockout medaka (the bottom row) compared with that of WT (the upper row). (D) The amino acid sequence of NPFF (pro-NPFF) or GnRH3 of knockout fish (the bottom row) compared with that of WT (the upper row). The changes in amino acid sequences caused by frame shift are indicated by blue font. * means stop codon. (E) Illustration of the side view of the fish brain and the frontal section in the telencephalon. The red line indicates the region of the section of F and G, and the red rectangular indicates the brain region around the terminal nerve. (F) NPFF-immunoreactive cell bodies were found only in the brain of WT. The black arrowhead shows cell bodies of TN-GnRH3 neurons. The dotted arrowhead indicates the region of TN-GnRH3 neurons. Scale bar: 20 µm. (G) GnRH3-immunoreactive cell bodies and fibers were found only in the brain of WT. The white arrowhead shows cell body of TN-GnRH3 neurons. The dotted arrowhead indicates the region of TN-GnRH3 neurons. † means non-specific signal of the meninges. Scale bar: 25 µm.

**Figure 1—figure supplement 1:**
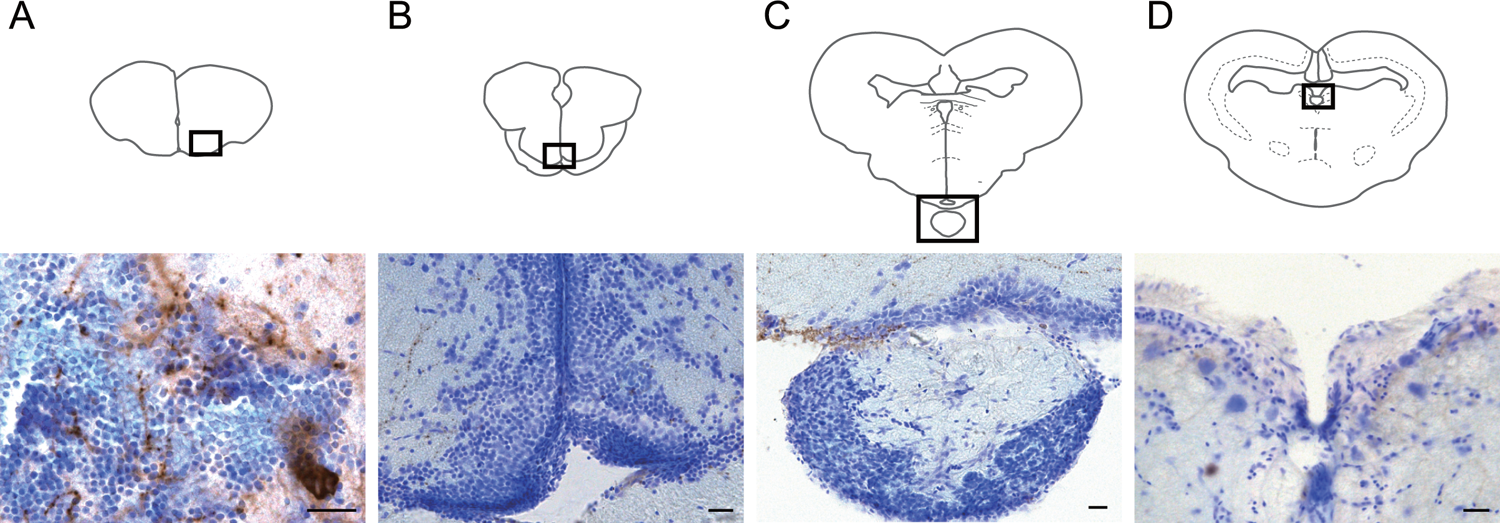
LRH13 is a specific antibody for medaka GnRH3 (A) LRH13 immuno-reactive cell bodies and neurites were localized in the terminal nerve. On the other hand, LRH13 immuno-reactive cell bodies were found neither in the POA (B), where the cell bodies of GnRH1 neurons are located, and LRH13 immuno-reactive fibers, nor in the pituitary (C), where the axons of GnRH1 neurons are located. In addition, immuno-reactive structures were not found in the midbrain tegmentum (D), where the cell bodies of GnRH2 neurons are localized. Scale bar: 20 µm.

In the present study, in order to examine how neuropeptide in TN-GnRH neurons modulate sexual behavior, we generated GnRH and/or NPFF knockout medaka, a teleost, and analyzed their sexual behaviors. Medaka offers various advantages for conducting the present study; medaka regularly shows stereotypical sexual behavior every day (Ono and Uematsu, 1957, Hiraki-Kajiyama et al., 2019). In addition, unlike mammals and zebrafish, medaka is one of the animals in which one of the three paralogues of GnRH, *gnrh3*, is clearly localized in the terminal nerve neurons in the brain. This makes it easy to analyze the functions of GnRH (GnRH3) in the terminal nerve independently of the hypophysiotropic GnRH (GnRH1) neurons (Okubo and Nagahama, 2008, Kim et al., 2011). Furthermore, in common with the previous report in the dwarf gourami (Saito et al., 2010), NPFF is expressed only in the TN-GnRH neurons in the medaka brain (Figure 1A).

Here, we performed behavioral analyses using *gnrh3* knockout (*gnrh3 ^-/-^*) and/or *npff* knockout (*npff ^-/-^*) medaka and found that NPFF and GnRH3 modulate male sexual behaviors. In addition, to localize the target neurons of NPFF or GnRH3 in the brain, we anatomically analyzed the expressions of NPFF and GnRH receptors. Our results demonstrated that balanced release of NPFF and GnRH3 from the terminal nerve may play an important role especially in the control of motivation for male sexual behavior.

## Result

### Generation of *npff ^-/-^* and *gnrh3 ^-/-^* medaka

For the analysis of the function of NPFF and GnRH3, we generated knockout medaka of each gene and double knockout medaka. Our designed TALEN caused frame shift of *npff* and *gnrh3* (Figure 1B and 1C) and change of amino acid in each core peptide (Figure 1D). In addition, GnRH3 and NPFF double knockout medaka (*gnrh3 ^-/-^*;*npff ^-/-^*) was generated by intercrossing *npff ^-/-^* medaka and *gnrh3 ^-/-^* medaka. Next, we examined whether TALEN induced null function.

Because NPFF in the terminal nerve can be labeled by anti-FMRF (Phe-Met-Arg-Phe-NH2) antibody (Saito et al., 2010), we made frontal brain sections of WT and *npff ^-/-^* (Figure 1E) and performed immunohistochemistry (IHC) using this antibody. The FMRF-immunoreactive signals were found in the cell bodies of TN-GnRH neurons in WT but not in *npff ^-/-^* medaka (Figure 1F). Considering the method for generation of *gnrh3 ^-/-^*;*npff ^-/-^* medaka, *gnrh3 ^-/-^*;*npff ^-/-^* medaka should also have lost NPFF expression. On the other hand, to examine the loss of GnRH3 expression, we performed IHC using anti-GnRH3 antibody (LRH13), which is a specific antibody for GnRH3 in medaka (Figure 1—figure supplement 1). As shown in Figure 1G, GnRH3-immunoreactive signals were found in WT brain but not in *gnrh3 ^-/-^* or *gnrh3 ^-/-^*;*npff ^-/-^* medaka. These results confirm that our knockout strains have lost the functional peptides of NPFF and/or GnRH3.

### *npff ^-/-^* male shows a delay in various sexual behavior repertories

To examine whether NPFF and/or GnRH3 modulate motivation for sexual behavior, we first analyzed sexual behaviors of *npff ^-/-^* medaka and compared those behaviors with WT ones (Figure 2). We recorded sexual behaviors of *npff ^-/-^* and WT medaka that have fully grown up (Figure 2— figure supplement 1). Sexual behaviors of *npff ^-/-^* medaka showed some abnormalities compared with WT. The stereotypical sexual behavior repertories of medaka are illustrated in Figure 2A. First, the male medaka follows or chases a female (called following) and presents courtship behavior to the female by quickly circling in front of the female (courtship). If the female accepts the male’s courtship, the male holds the female’s body by its anal fin (clasping), which is followed by simultaneous spawning (spawning). Here, we found that *npff ^-/-^* males show a prolongation of the latency to the first following, the first courtship, the first clasping, and spawning (Figure 2B-E). In addition, in the pair of *npff ^-/-^* male and WT female, the courtship frequency before spawning (Figure 2—figure supplement 2A) and percentage of following with successful courtship (Figure 2—figure supplement 2B) were also significantly less than those in WT pairs. On the other hand, *npff ^-/-^* female showed a prolongation of the latency to clasping and spawning (Figure 2D and 2E). It should be noted that we used all pairs that spawned at least before the experimental day. Considering these results, *npff ^-/-^* medaka, especially *npff ^-/-^* male, are suggested to have less motivation for sexual behaviors (see Discussion for detail).

**Figure 2:**
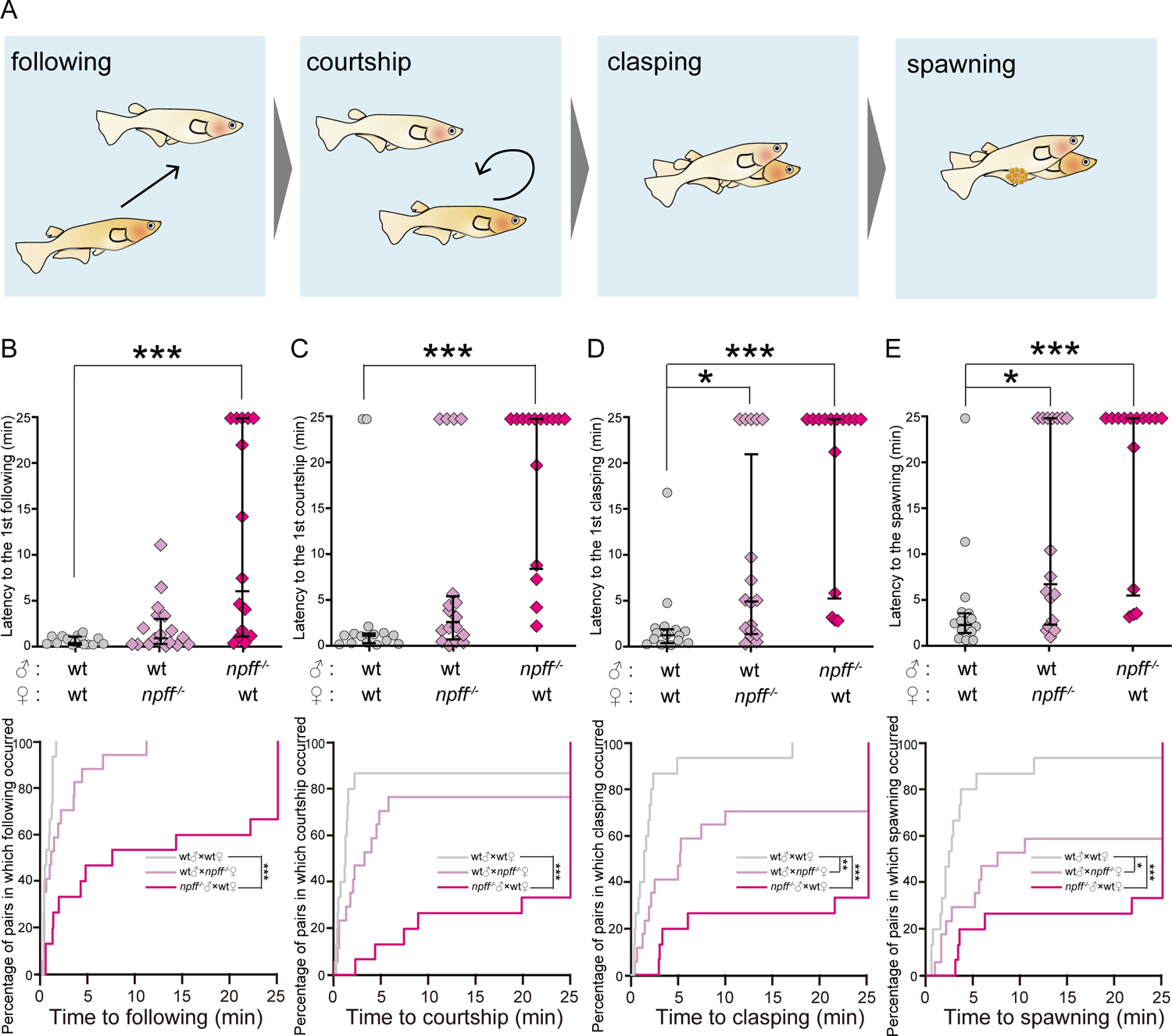
*npff ^-/-^* medaka shows the increase in the latency to sexual behaviors (A) Normal sexual behavioral repertories in WT male and female medaka. (B-E) The upper graph: latency to each sexual behavior illustrated in (A). The lower graph: reverse Kaplan-Meier curve indicating the percentage of pairs in which each sexual behavior illustrated in (A) occurred. The short latency indicates that the sequence of sexual behavior proceeded quickly. The latency of a pair, which did not show the behavior, was defined as 25 min. (B) The first following. (C) The first courtship. (D) The first clasping. (E) The spawning. WT male and female pair: n = 15, WT male and *npff ^-/-^* female pair: n = 17, *npff ^-/-^* male and WT female pair: n =15. For the analyses of the latency, each knockout group was compared with WT male and female pair (*: p < 0.05, **: p < 0.01, ***: p < 0.001, Steel’s multiple comparison test). For the analyses of percentage of pairs, we used Log-rank test with Bonferroni’s correction (*: p < 0.05, **: p < 0.01, ***: p < 0.001).

**Figure 2—figure supplement 1:**
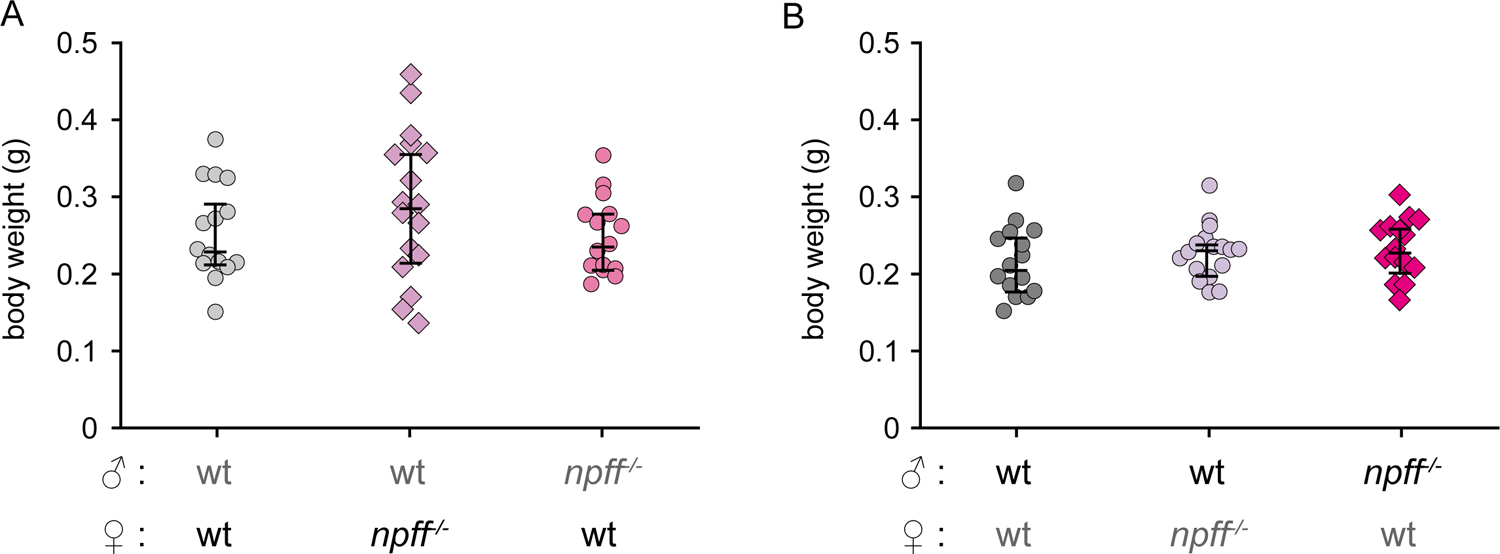
*npff ^-/-^* medaka shows normal body size All knockout fish grew as normally as WT. (A) & (B) Body weight of control (WT) and *npff ^-/-^* medaka. WT male and female pair: n = 15, WT male and *npff ^-/-^* female pair: n = 17, *npff ^-/-^* male and WT female pair: n = 15. Steel’s multiple comparison test, not significant. The body weights of the fish labeled with bold font are shown. The fish labeled with gray font indicates that of the partner in behavioral analysis.

**Figure 2—figure supplement 2:**
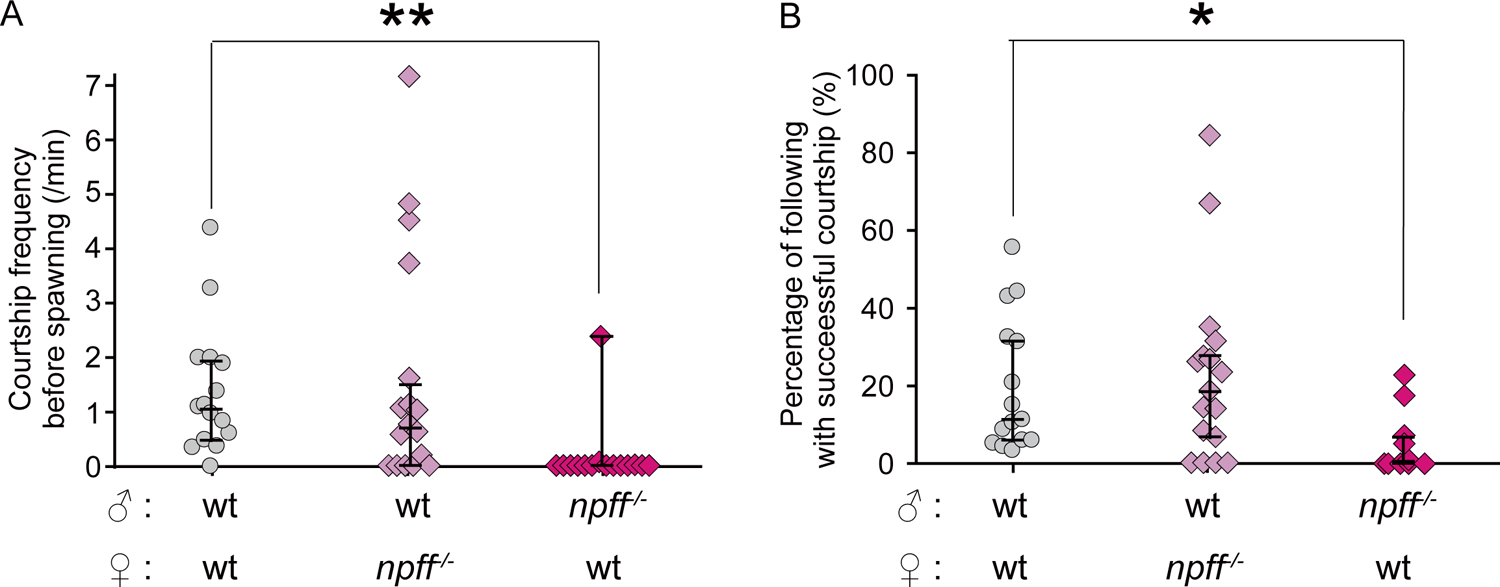
*npff ^-/-^* male shows an impairment of following and courtship behaviors (A) courtship frequency before spawning (times/min). WT male and female pair: n = 15, WT male and *npff ^-/-^* female pair: n = 17, *npff ^-/-^* male and WT female pair: n = 15. (B) percentage of following with successful courtship (%). WT male and female pair: n = 15, WT male and *npff ^-/-^* female pair: n = 17, *npff ^-/-^* male and WT female pair: n = 10. Each knockout group was compared with WT male and female pair. (*: p < 0.05, **: p < 0.01, Steel’s multiple comparison test)

### *gnrh3 ^-/-^* male shows a delay in sexual behavior repertories after courtship

Next, to examine the function(s) of GnRH3 in the control of sexual behavior, we conducted behavioral analysis of *gnrh3 ^-/-^* or *gnrh3 ^-/-^*;*npff ^-/-^* (Figure 3) in addition to *npff ^-/-^* (Figure 2). We recorded sexual behaviors of each pair of medaka that have fully grown up (Figure 3—figure supplement 1). Then, in the pairs of *gnrh3 ^-/-^* male and WT female, the latencies to the first clasping and spawning were significantly longer than WT pairs (Figure 3B-E). Similar to the experiments of *npff ^-/-^*, we used all pairs that spawned at least before the experimental day. In addition, because there was no significant difference in the courtship frequency before spawning (Figure 3—figure supplement 2A) and percentage of following with successful courtship (Figure 3—figure supplement 2B) among all the pairs, it is suggested that *gnrh3 ^-/-^* male are not able to show sexual behaviors smoothly after courtship. On the other hand, *gnrh3 ^-/-^*;*npff ^-/-^* male and female showed normal sexual behaviors (Figure 3B-E).

**Figure 3:**
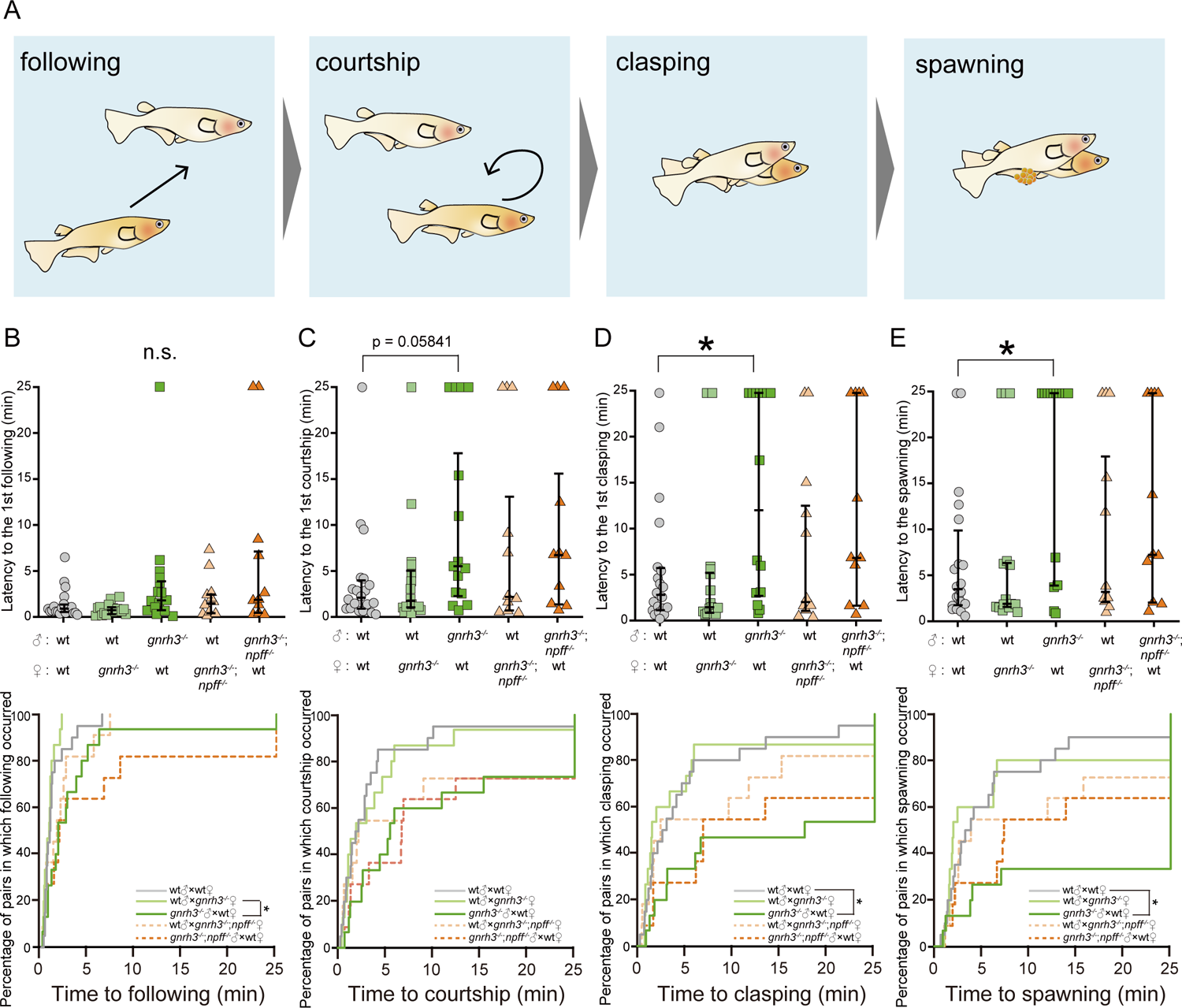
*gnrh3 ^-/-^* male shows the increase in the latency to some sexual behaviors, although *gnrh3 ^-/-^*;*npff ^-/-^* male shows normal sexual behaviors (A) Normal sexual behavioral repertories in WT male and female medaka. (B-E) The upper graph: latency to each sexual behavior illustrated in (A). The lower graph: reverse Kaplan-Meier curve indicating the percentage of pairs in which each sexual behavior illustrated in (A) occurred. The short latency indicates that the sequence of sexual behavior proceeded quickly. The latency of a pair, which did not show the behavior, was defined as 25 min. (B) The first following. (C) The first courtship. (D) The first clasping. (E) The spawning. WT male and female pair: n = 20, WT male and *gnrh3 ^-/-^* female pair: n = 15, *gnrh3 ^-/-^* male and WT female pair: n = 15, WT male and *gnrh3 ^-/-^*;*npff ^-/-^* female pair: n = 11, *gnrh3 ^-/-^*;*npff ^-/-^* male and WT female pair: n = 11. For the analyses of the latency, each knockout group was compared with WT male and female pair (*: p < 0.05, Steel’s multiple comparison test). For the analyses of percentage of pairs, we used Log-rank test with Bonferroni’s correction (*: p < 0.05).

**Figure 3—figure supplement 1:**
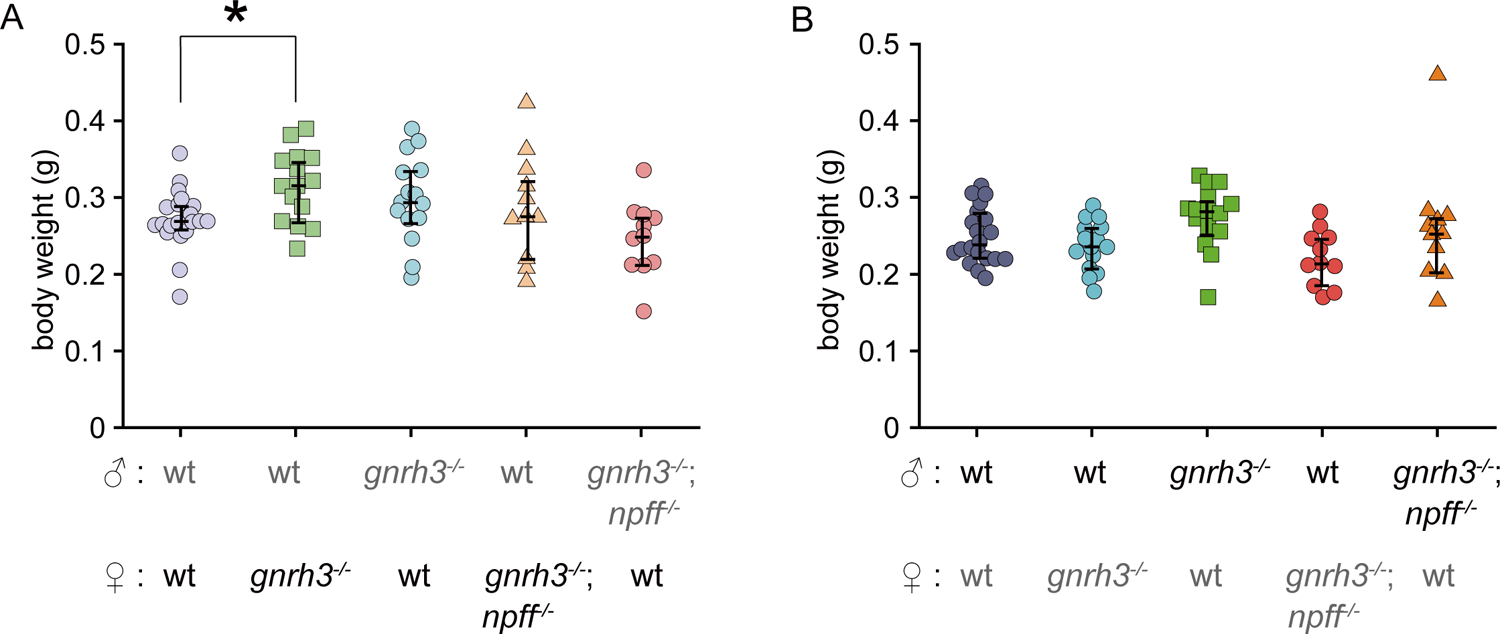
*gnrh3 ^-/-^* and *gnrh3 ^-/-^*;*npff ^-/-^* medaka showed normal body size All knockout fish grew almost normally as well as WT. (A) & (B) Body weight of control (WT), *gnrh3 ^-/-^*, and *gnrh3 ^-/-^*;*npff ^-/-^* medaka. WT male and female pair: n = 20, WT male and *gnrh3 ^-/-^* female pair: n = 15, *gnrh3 ^-/-^* male and WT female pair: n = 15, WT male and *gnrh3 ^-/-^*;*npff ^-/-^* female pair: n = 11, *gnrh3 ^-/-^*;*npff ^-/-^* male and WT female pair: n = 11. Steel’s multiple comparison test, *: p < 0.05. The body weights of the fish labeled with bold font are shown. The fish labeled with gray font indicates that of the partner in behavioral analysis.

**Figure 3—figure supplement 2:**
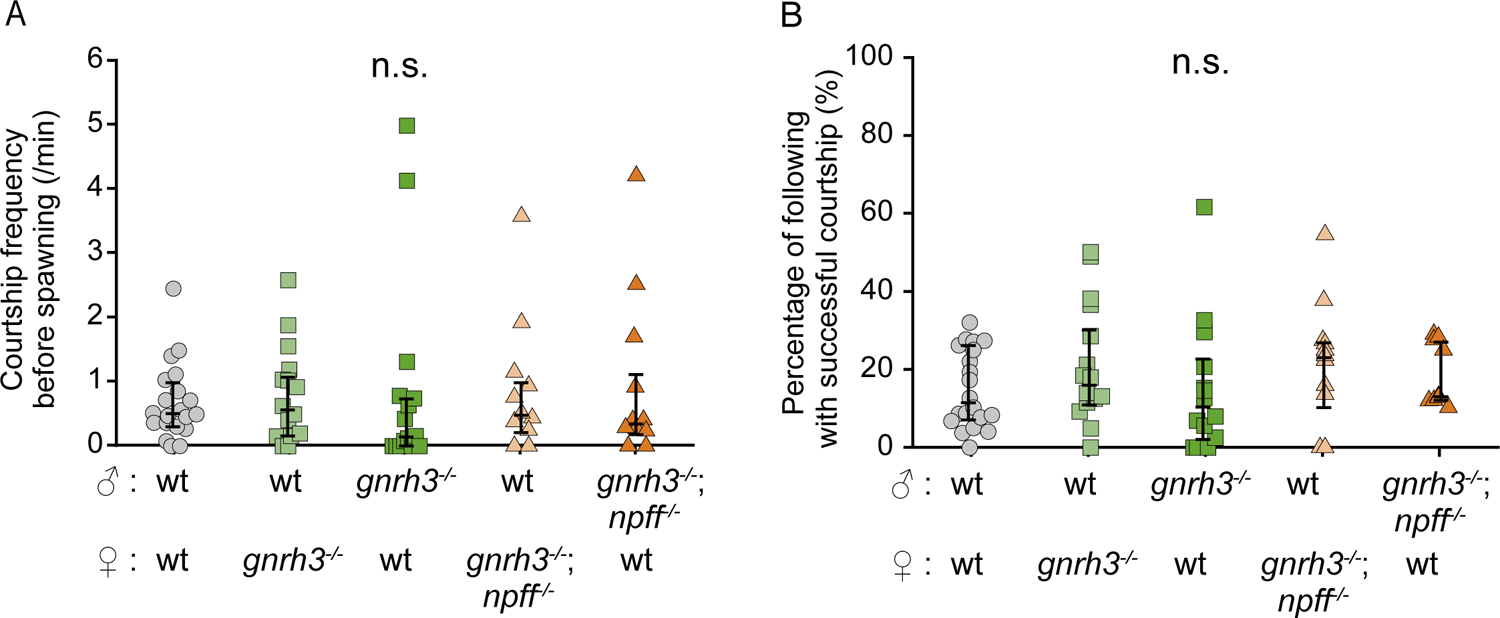
*gnrh3 ^-/-^* and *gnrh3 ^-/-^*;*npff ^-/-^* male show the normal following and courtship behaviors (A) Courtship frequency before spawning (times/min). WT male and female pair: n = 20, WT male and *gnrh3 ^-/-^* female pair: n = 15, *gnrh3 ^-/-^* male and WT female pair: n = 15, WT male and *gnrh3 ^-/-^*;*npff ^-/-^* female pair: n = 11, *gnrh3 ^-/-^*;*npff ^-/-^* male and WT female pair: n = 11. (D) Percentage of following with successful courtship (%). WT male and female pair: n = 20, WT male and *gnrh3 ^-/-^* female pair: n = 15, *gnrh3 ^-/-^* male and WT female pair: n = 14, WT male and *gnrh3 ^-/-^*;*npff ^-/-^* female pair: n = 11, *gnrh3 ^-/-^*;*npff ^-/-^* male and WT female pair: n = 9. Each knockout group was compared with WT male and female pair. (n.s.: not significant, Steel’s multiple comparison test)

### NPFF receptors are expressed broadly in the brain

Since *npff ^-/-^* showed abnormal sexual behaviors, we examined the possible neuronal circuit via NPFF, which is expressed only in TN-GnRH neurons in the brain, by analyzing anatomical localizations of NPFF receptors in the brain. First, by the 5’ and 3’ rapid amplification of cDNA ends (RACE), we found three genes as NPFF receptors and identified *gpr147* (so-called NPFFR1, Bonini et al., 2000), *gpr74-1* and *gpr74-2* (so-called NPFFR2, Bonini et al., 2000) (Figure 4— figure supplement 1). All of these receptors were shown to function as G-protein coupled receptors, whose ligand is NPFF (Figure 4—figure supplement 2). Then, by *in situ* hybridization of these genes, we found that these receptors are expressed broadly in the brain (Figure 4A, abbreviation of brain region in Supplementary file 1). Because TN-GnRH neurons do not project to the pituitary, we analyzed brain regions except the pituitary. In Vv and Vs, which are the brain region involved in the control of sexual behavior (Koyama et al., 1984, Satou et al., 1984), *gpr74-2* and *gpr147* were abundantly expressed, respectively (Figure 4B and 4C, abbreviation of brain nuclei is described in Supplementary file 1). In addition, in POA and hypothalamus, all subtypes of NPFF receptors were expressed (Figure 4D). *gpr74-1* was expressed in the dorsal and ventral part of POA, while *gpr74-2* and *gpr147* were mostly expressed in the dorsal part of POA. Furthermore, TN-GnRH neurons, which have been suggested to modulate the motivation for sexual behavior (Wirsig and Leonard, 1987, Yamamoto et al., 1997), also expressed *gpr74-1* (Figure 4E). Thus, it is also suggested morphologically that NPFF plays an important role in sexual behaviors.

**Figure 4:**
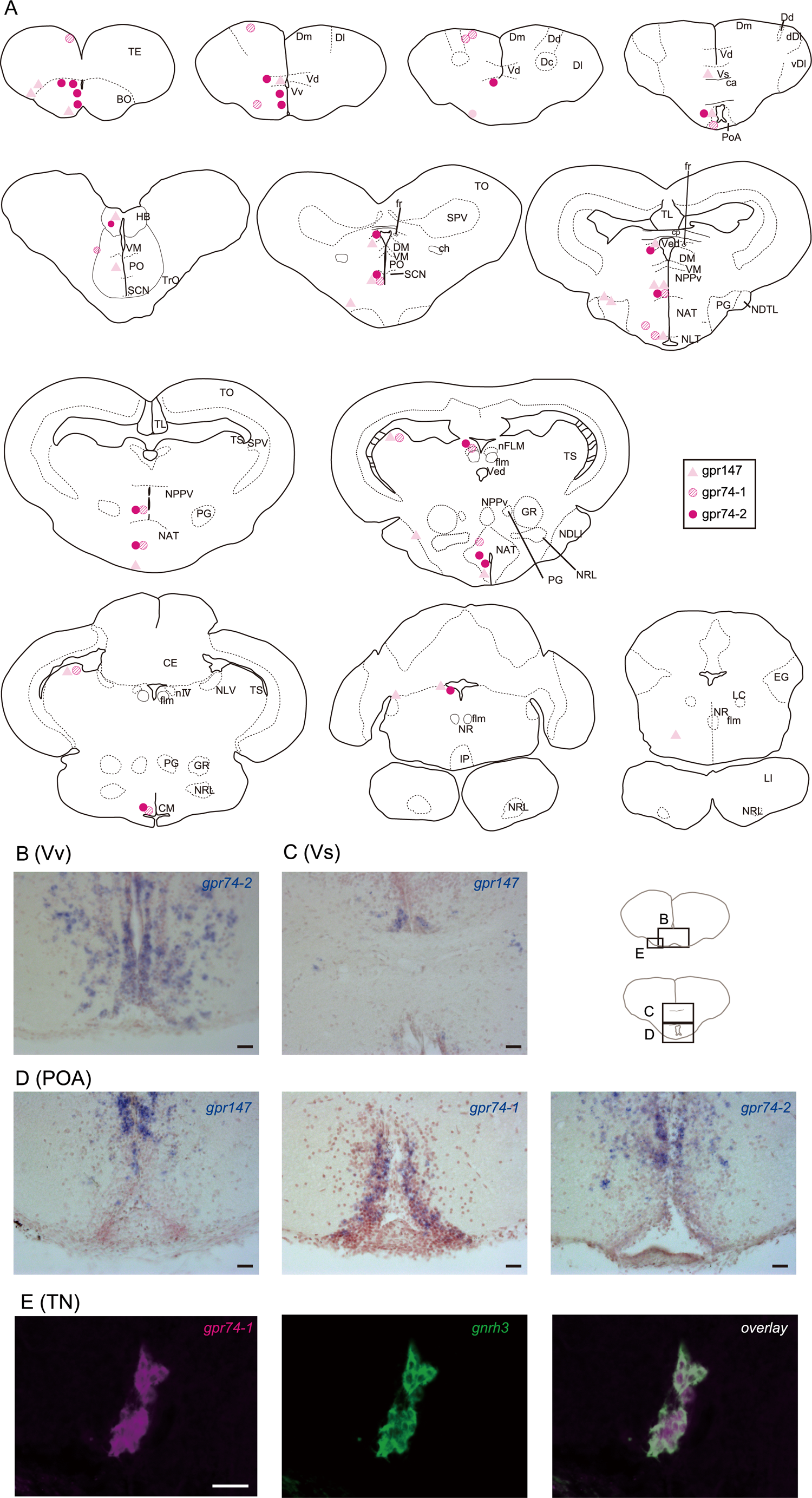
*gpr147*, *74-1*, and *74-2* (NPFF receptors)-expressing neurons are distributed broadly in the brain. (A) Localization of the cell bodies expressing *gpr147*, *74-1*, and *74-2*. The right part of the brain indicates nuclear boundaries, while the left part of the brain does localizations of neurons expressing each type of receptor. (B) *gpr74-2* expressing neurons in the ventralis telencephali pars ventralis (Vv) (C) *gpr147* expressing neurons in the ventralis telencephali pars supracommissuralis (Vs) (D) *gpr147*, *74-1*, and *74-2* expressing neurons in the preoptic area (POA) (E) Co-expression of *gpr74-1* (magenta) and *gnrh3* (green) in the terminal nerve. Scale bar: 20 μm.

**Figure 4—figure supplement 1:**
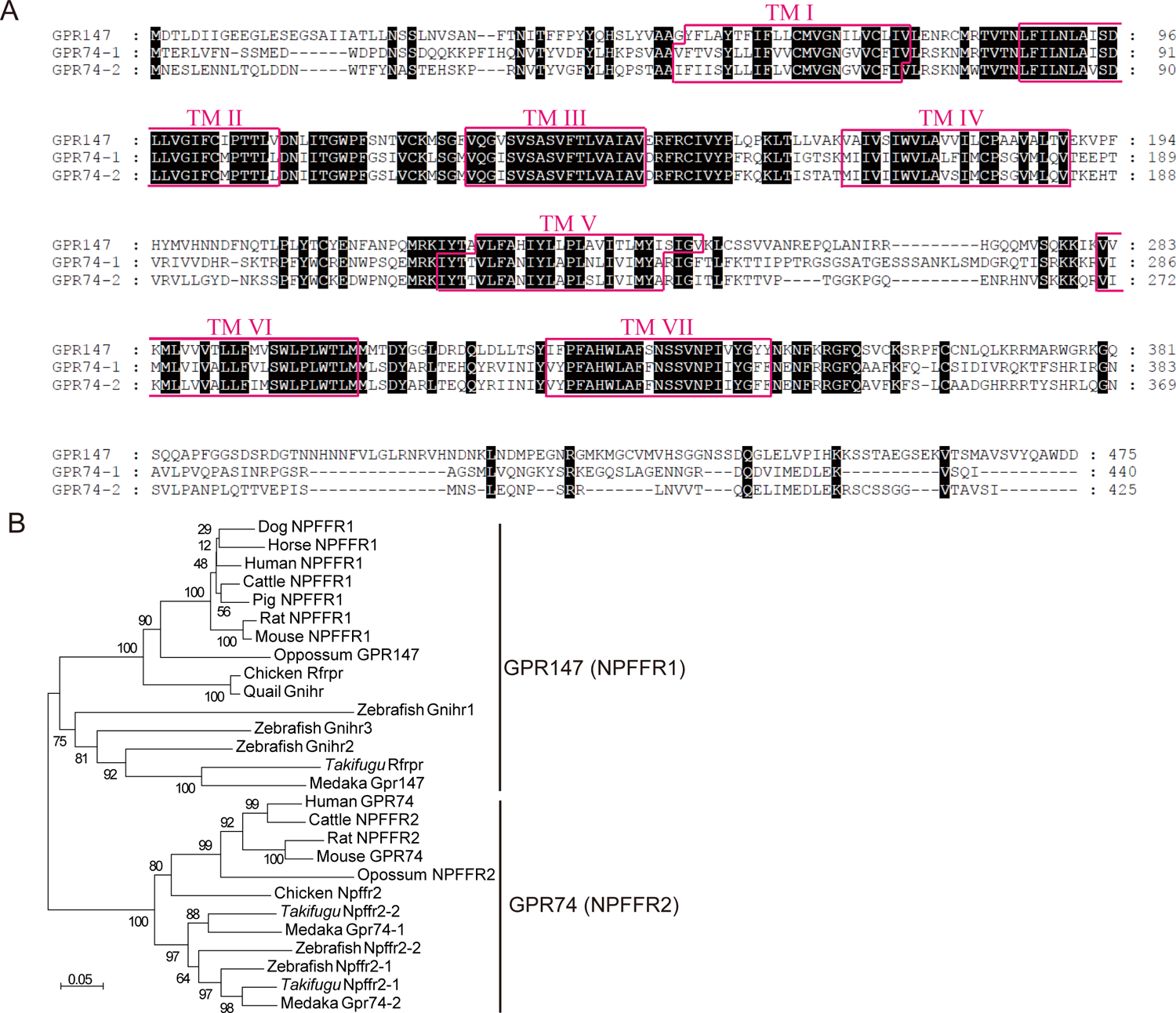
Alignment of the deduced amino acid sequence and phylogenic tree of Gpr147, 74-1, and 74-2 (A) The amino acid sequence of Gpr147, 74-1, and 74-2. Full-length protein sequences were aligned by ClustalX 2.1 using the default settings. Identical residues are shaded in black. (B) Phylogenetic tree of GPR147, GPR74-1, and GPR74-2. Unrooted phylogenetic tree of the receptors was constructed by the neighbor-joining method using MEGA6 software. Each medaka receptor was named Gpr147, Gpr74-1, or Gpr74-2 in accordance with other known sequences of GPR147 and GPR74. The numbers at the nodes are bootstrap values as percentages, which were obtained from 1,000 replicates. Scale bar indicates the substitution rate per residue.

**Figure 4—figure supplement 2:**
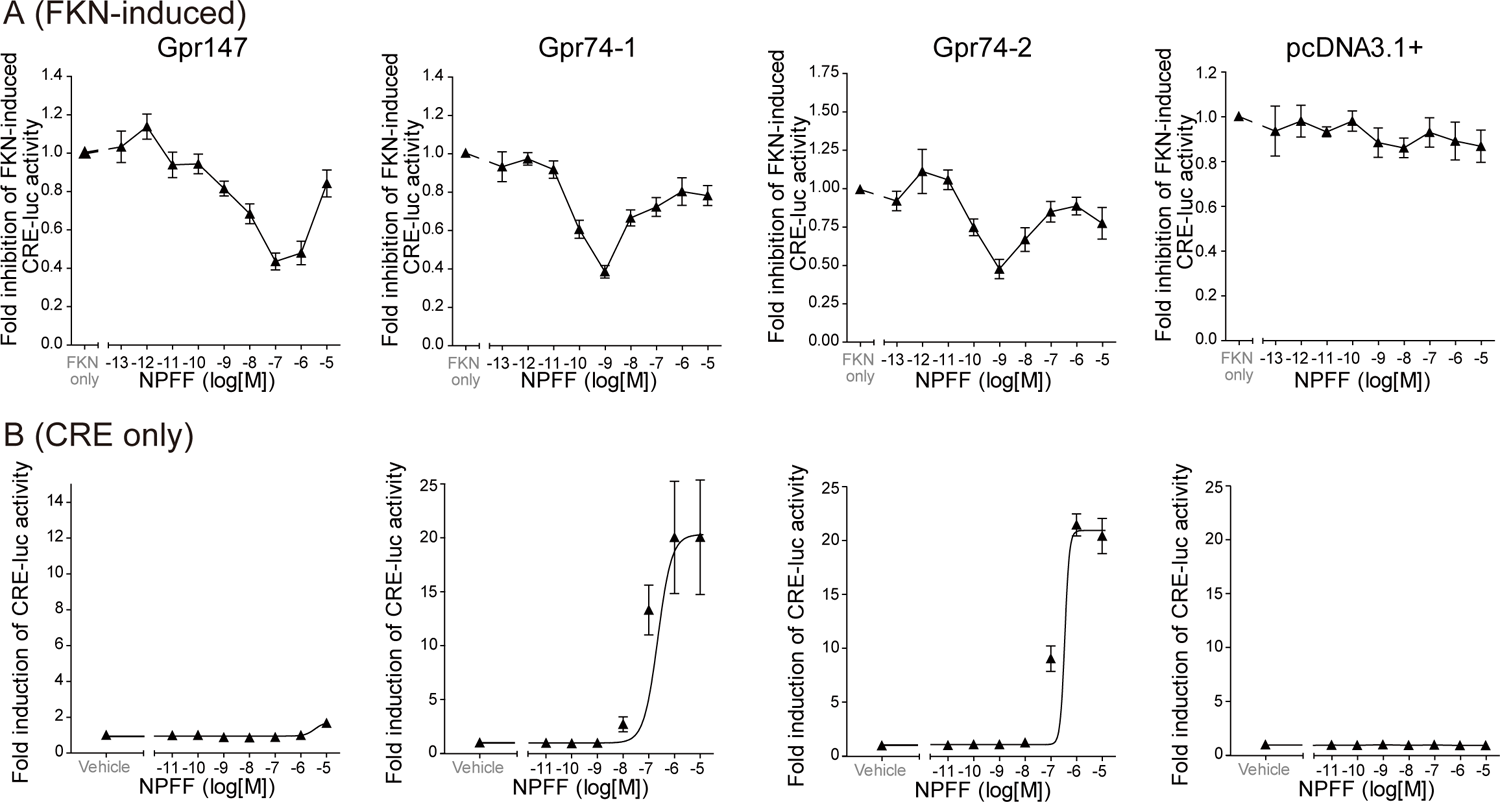
Cloned Gpr147, 74-1, and 74-2 worked as G-protein coupled receptor. (A) Inhibition of forskolin-induced CRE-luciferase activities in HEK293T cells transfected with medaka Gpr147, Gpr74-1 or Gpr74-2 in combination with pGL4.29 (CRE-luc) plasmid. The mammalian expression vector pcDNA3.1+ was transfected as a mock-transfection. (D) Induction of CRE-luciferase activities HEK293T cells transfected with medaka Gpr147, Gpr74-1 or Gpr74-2 in combination with pGL4.29 (CRE-luc) plasmid. The mammalian expression vector pcDNA3.1+ was transfected as a mock-transfection. In A and B, after stimulation with increasing concentration of medaka Npff (dark triangle) luciferase activity was assessed. The data are indicated as the means ± S.E.M. of at least three independent experiments performed in triplicate.

### GnRH receptors are mainly expressed in the sensory processing areas of the brain

We next examined the possible neuronal circuit via GnRH3 by analyzing anatomical localizations of GnRH receptors in the brain (Figure 5A, abbreviation of brain region in Supplementary file 1). Medaka has four subtypes of GnRH receptors (we here followed the terminology of Okubo et al., 2003, Kim et al., 2011; GnRHR1-4). *gnrhr1* (Figure 5B) and *gnrhr2* (Figure 5C) were expressed in two different brain regions. *gnrhr1* was expressed in the Dp and the NI. There were more neurons expressing *gnrhr1* in the Dp than in the NI. *gnrhr2* was expressed in the pineal organ and the NDTL. *gnrhr2*-expressing neurons in the pineal organ were located in the surface of the organ. *gnrhr3* (Figure 5D), among the four subtypes, was most broadly expressed in the brain: Vv, Dd, vDm, and POA in the telencephalon; PO, NAT, NVT, and NRP in the hypothalamus; MCN and near flm in the medulla oblongata. Since *gnrhr4* in medaka was newly discovered in 2011 (Kim et al., 2011), we examined whether GnRHR4 functions as a Gq-coupled receptor and found that it does so (Figure 5—figure supplement 1). *gnrhr4* (Figure 5E) was expressed in the NAT and the NFLM. *gnrhr4*-expressing neurons were localized in NFLM as a symmetrical pair.

**Figure 5:**
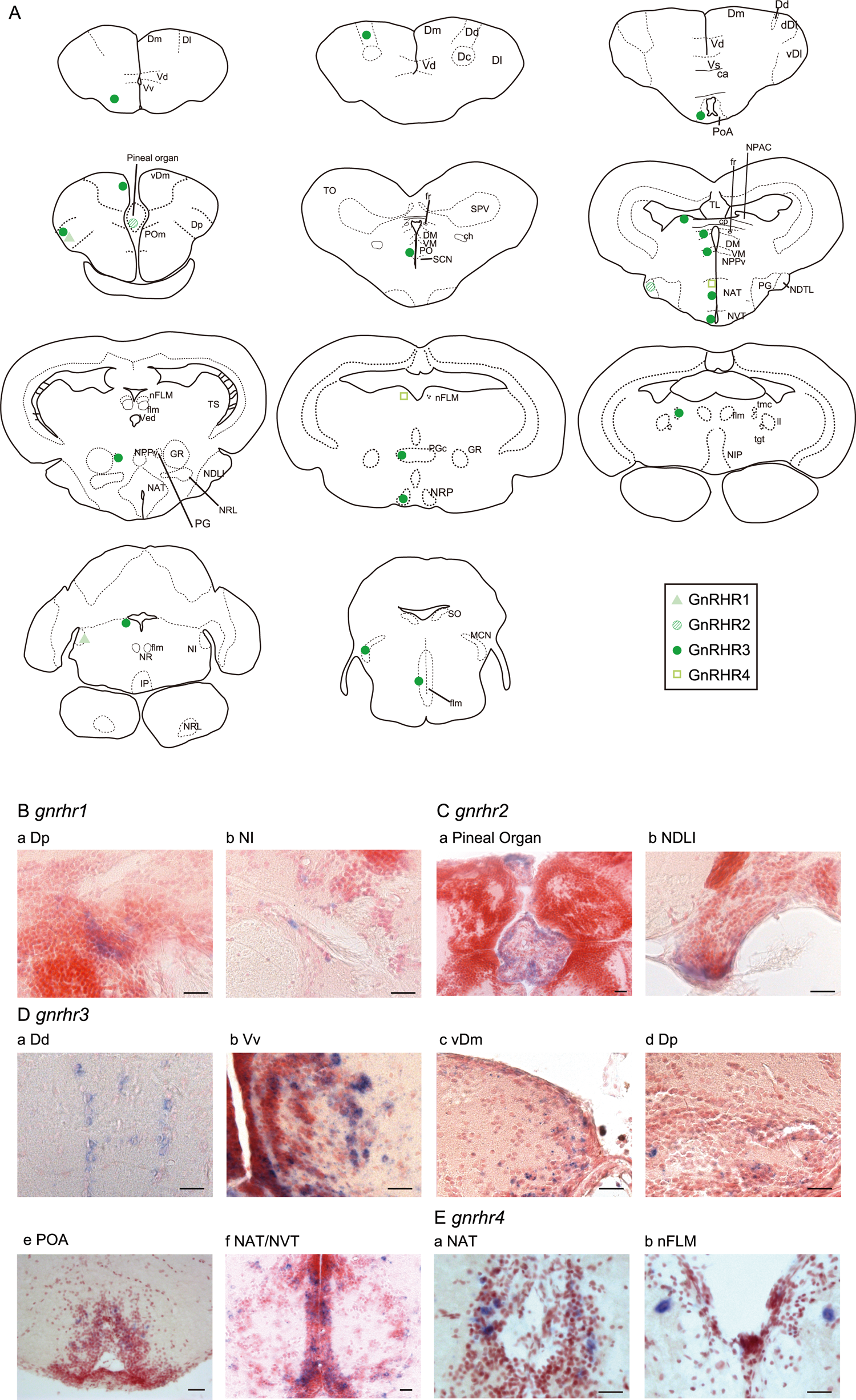
*gnrhr1*, *2*, and *4* -expressing neurons are distributed locally, whereas *gnrhr3*-expressing neurons are distributed broadly in the brain (A) Localization of the cell bodies expressing each *gnrhr*. The right part of the brain indicates nuclear boundaries, while the left part of the brain does localizations of neurons expressing each type of receptor. (B) *gnrhr1*-expressing neurons in the area dorsalis telencephali pars posterior (A-a) and the nucleus isthmi (A-b). (C) *gnrhr2*-expressing neurons in the pineal organ (C-a) and the nucleus diffusus tori lateralis (C-b). (D) *gnrhr3-*expressing neurons in representative brain nuclei. a, area dorsalis telencephali pars dorsalis (Dd), b, area ventralis telencephali pars ventralis (Vv), c, ventral region of Dm (vDm), d, area dorsalis telencephali pars posterior, e, preoptic area (POA), f, nucleus anterior tuberis (NAT) and neucleus ventralis tuberis (NVT), (E) *gnrhr4* was expressed in the nucleus anterior tuberis (E-a) and the nucleus of fasciculus longitudinalis mediali (E-b). scale bar in B-E: 20 μm.

**Figure 5—figure supplement 1:**
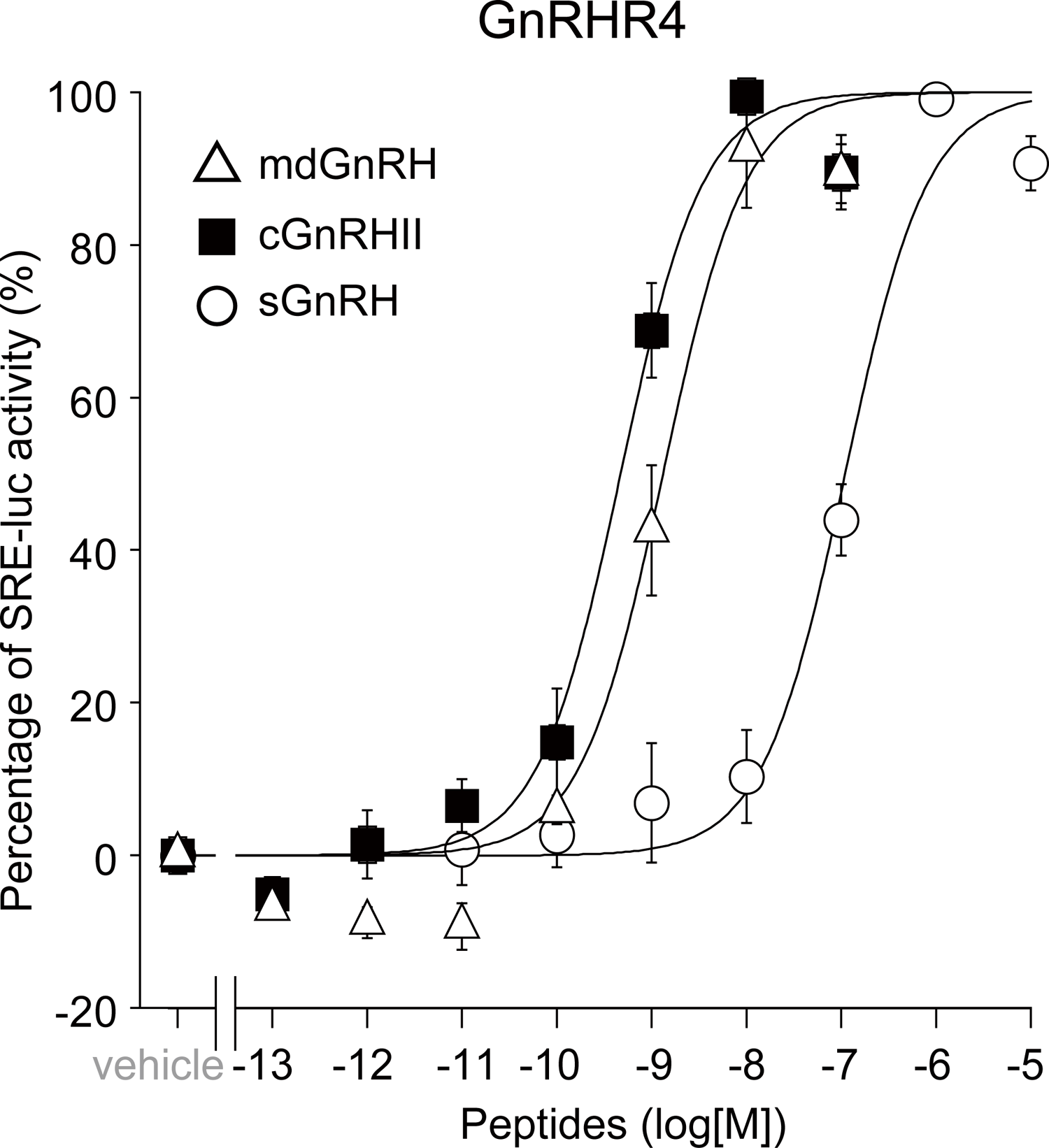
GnRHR4 functions as a Gq -protein coupled receptor. Induction of SRE-luciferase activities HEK293T cells transfected with medaka *gnrhr4* in combination with pGL4.33 (SRE-luc) plasmid. After stimulation with increasing concentration of medaka GnRH1 (mdGnRH, white triangle), chicken GnRH2 (cGnRHII, dark square), or salmon GnRH3 (sGnRH, white circle), luciferase activity was assessed. The data are indicated as the means ± S.E.M. of at least three independent experiments performed in triplicate.

### The expression level of *immediate early gene*, *egr1*, in vPOA of *npff ^-/-^* male is lower than that of WT

As described above, *npff ^-/-^* male showed severe delay in sexual behaviors. To examine the neural activities in WT and *npff ^-/-^* male, we compared the expression of one of the immediate early genes, *egr1*, in *npff ^-/-^* male with that in WT male. *egr1* expression is suggested to represent neuronal activities during 30 min before sacrifice (Burmeister and Fernald, 2005). We made pairs according to the behavioral test of *npff ^-/-^* described above, sacrificed the male, and dissected out the brains 25 – 28 min after removing the transparent separation, when the motivation for sexual behavior in WT male was observed to be high in the present behavioral analysis (see Figure 2). Because WT showed more *egr1*-expessing neurons only in the ventral part of preoptic area (vPOA) than *npff ^-/-^* male among other brain regions, we examined co-expression of *egr1* and *gpr74-1,* which is the *npff receptor* gene expressed in the vPOA (Figure 4D). As shown in Figure 6A and 6B, the percentage of *egr1*-expessing neurons in the vPOA of *npff ^-/-^* male was less than that of WT, while the difference between the percentage of *gpr74-1*-expessing neurons in *npff ^-/-^* and that in WT was not significant (Figure 6C). On the other hand, we found a small number of *egr1* and *gpr74-1* co-expressing neurons in both genotypes (WT: 30 ± 8 cells, *npff ^-/-^*: 13 ± 7 cells), although the difference in the percentage of *egr1*- and *gpr74-1*-coexpessing neurons was not significant between the two (Figure 6D). It should be noted that all WT males spawned within 6 min after removing the transparent separation, while five out of six *npff ^-/-^* males failed to spawn for 25 min after removing the transparent separation. Taken together, it is suggested that NPFF activates neurons in the vPOA via NPFF receptor-expressing neurons, possibly via indirect pathway.

**Figure 6.**
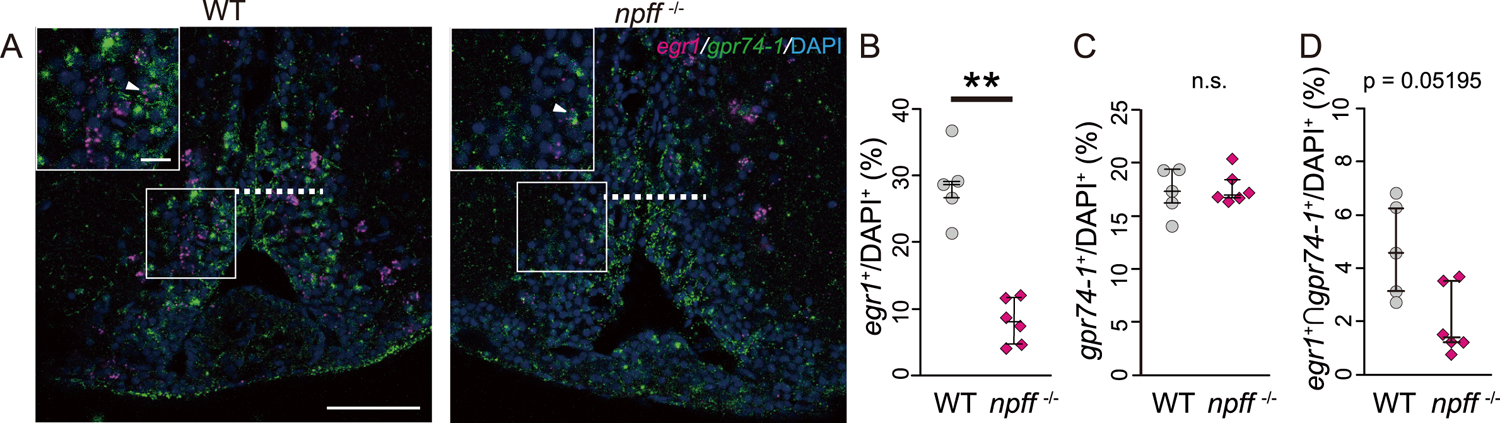
: e*gr1*-expressing neurons were more abundantly observed in WT than in *npff ^-/^* male medaka (A) *egr1* (magenta) and *gpr74-1* (green) expression in the preoptic area (POA) of WT (left) and *npff ^-/-^* (right) male medaka. The nuclei were counterstained by DAPI (blue). The inset shows an enlarged image of the region in square. The white arrowhead indicates a neuron expressing both *egr1* and *gpr74-1*. We counted cells in vPOA (the region under dotted bars). Scale bars: 50 μm, 10 μm (inset) (B) The percentage of *egr1-*expressing neurons (*egr1^+^*) in the ventral part of POA. (C) The percentage of *gpr74-1^+^-*expressing neurons (*gpr74-1^+^*) in the ventral POA. (D) The percentage of neurons expressing both *gpr74-1* and *egr1* (*egr1^+^*∩*gpr74-1^+^*) in the ventral POA. WT: n = 5, *npff ^-/-^*: n = 6. Mann-Whitney U test, **: p < 0.01, n.s.: not significant.

### The expression level of *npff* in *gnrh3 ^-/-^* medaka is lower than that in WT medaka

Our behavioral analyses suggested that NPFF and GnRH3 may facilitate the motivation for the control of male sexual behaviors in slightly different manners. Therefore, to examine the relationship of GnRH3 and NPFF, we analyzed the expression of *npff* or *gnrh3* in *npff ^-/-^*, *gnrh3^-/-^*, and WT medaka. As shown in Figure 7A, the expression level of *npff* in *npff ^-/-^* was lower than that in WT, and that of *gnrh3* in *gnrh3 ^-/-^* was lower than that in WT, which suggests that nonsense-mediated mRNA decay has actually occurred. On the other hand, the expression level of *npff* in *gnrh3 ^-/-^* was lower than that in WT, while *npff ^-/-^* showed normal expression of *gnrh3*. Thus, it is suggested that GnRH3 may regulate the expression of *npff*.

**Figure 7.**
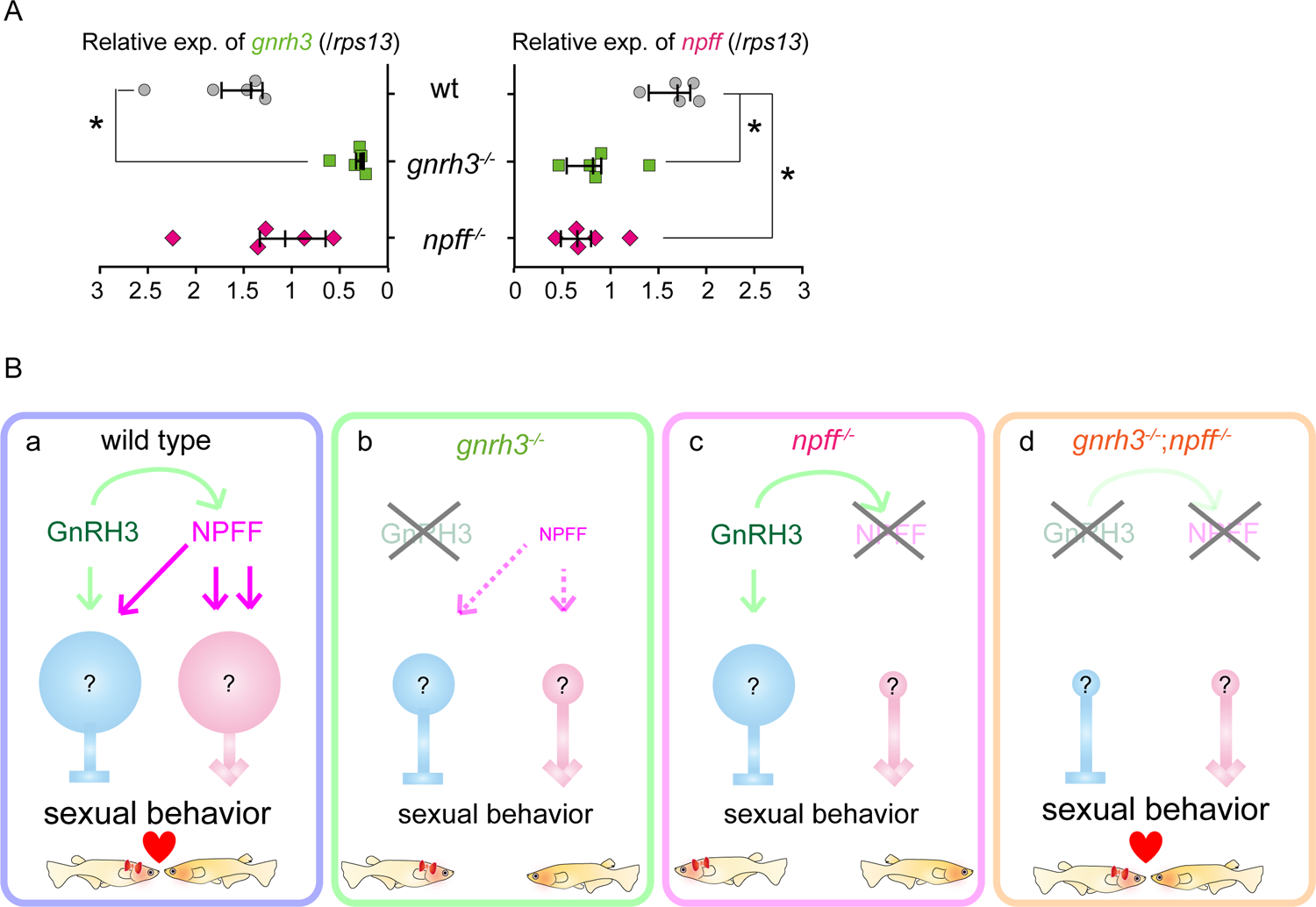
: Expression of *npff* in *gnrh3^-/-^* male is lower than that of WT male medaka (A) *gnrh3* and *npff* expression in the brain of *gnrh3 ^-/-^*, *npff ^-/-^*, and WT male medaka. Each group: n = 5. Steel’s multiple comparison test, *: p < 0.05. (B) Hypothetical illustrations of NPFF- and GnRH3-induced modulation of male sexual behavior. The size of the circle (blue or pink) indicates the activity level of each neural circuit. GnRH3 facilitates NPFF expression and activates inhibitory neural circuits (blue). On the other hand, NPFF activates both inhibitory (blue) and excitatory neural circuits (pink) controlling male sexual behaviors. (a) In WT, normal release of NPFF and GnRH3 causes balanced activities of excitatory and inhibitory neural circuits, which enables WT male to show normal sexual behaviors. (b) In *gnrh3^-/-^* medaka, loss of *gnrh3* expression and subsequently diminished expression of *npff* results in the low motivation for sexual behaviors. (c) In *npff ^-/-^*, loss of *npff* expression induces dominant activity of inhibitory neural circuits and diminishes motivation for sexual behaviors. (d) On the other hand, in *gnrh3 ^-/-^*;*npff ^-/^*^-^ male, loss of both genes results in the balanced activity of inhibitory and excitatory neural circuits, which leads to the normal sexual behaviors.

## Discussion

In the present study, we investigated the modulatory mechanism of sexual behaviors by TN-GnRH neurons. Results of the behavioral analyses of *gnrh3 ^-/-^* and/or *npff ^-/-^* medaka suggested that NPFF and GnRH3, which are co-expressed in TN-GnRH neurons, facilitate the motivation for male sexual behavior (Figure 2 and 3). In addition, our morphological analyses showed that NPFF receptors and GnRH receptors are expressed in the brain regions involved in sensory processing and control of sexual behavior (Figure 4 and 5). Furthermore, we demonstrated that NPFF activates neurons in the preoptic area via indirect pathway, which is considered to induce the increase in the motivation for male sexual behaviors (Figure 6). Since TN-GnRH neurons receive multiple sensory information, the present findings should provide novel knowledge for the study of sexual behaviors controlled by integration of multiple sensory inputs. Here, we discuss the modulatory mechanism of sexual behavior by NPFF and GnRH3.

### NPFF and GnRH3 modulate male sexual behaviors

First, the present behavioral study demonstrated that NPFF, which is expressed only in the TN-GnRH neurons in the brain, facilitates the motivation for sexual behaviors. We showed that the latency of all sexual behavioral repertories in *npff ^-/-^* male was longer than WT (Figure 2). The frequency of courtship before spawning, and the percentage of following with successful courtship in *npff ^-/-^* male was also less than WT (Figure 2—figure supplement 2). Since we could confirm that all pairs used in the present study spawned at least before the experimental day, it follows that all the sexual behavior repertories of *npff ^-/-^* male were delayed, but not disrupted after all. It should be noted that the duration of the analysis was set to 25 min, and the latency of the pair that did not show sexual behavior at all was defined as 25 min. Taken together, the present results clearly demonstrated that *npff ^-/-^* male shows severely low motivation for sexual behaviors. In other words, NPFF facilitates male sexual behaviors by modulating the motivational level. In addition, *npff ^-/-^* female showed the delay in clasping and spawning, which suggests that NPFF also facilitates the female acceptance of the male courtship behavior. To our knowledge, the present study is the first report of the function of NPFF in the control of sexual behaviors.

On the other hand, *gnrh3 ^-/-^* male showed a delay in clasping and spawning (Figure 3D and 3E). Since the frequency of courtship before spawning and the percentage of following with successful courtship in *gnrh3 ^-/-^* male were not significantly different from those in WT, following and courtship of *gnrh3 ^-/-^* male may be normal. These results suggest that *gnrh3 ^-/-^* male shows an impairment in the behavioral shift from courtship to clasping. For clasping, it has been suggested that male has to precisely recognize the distance to the female, the direction of female, the response of female to male courtship, etc. (Ono and Uematsu, 1957, Walter and Hamilton, 1970). Since TN-GnRH neurons are suggested to receive sensory information such as vision and olfaction (Yamamoto and Ito, 2000, Li et al., 2017), it is possible that *gnrh3 ^-/-^* male shows an impairment in the GnRH3-induced neuromodulation (see Umatani et al., 2015, Unatani and Oka, 2019) and cannot perform sexual behaviors smoothly. Unlike *gnrh3 ^-/-^* male, *gnrh3 ^-/-^* female showed normal sexual behaviors in the present study, although previous study reported that GnRH3 are involved in the female preference for the familiarized male (Okuyama et al., 2014). While the previous study focused on the social behavior by analyzing female preferences with two males, our study focused more on the instinctive sexual behavior between a pair of one male and one female. This difference in the analysis may have caused the apparent difference in phenotype of *gnrh3^-/-^* female. It is possible that, under some specific conditions where female has to actively choose male, GnRH3 may play an important role in the female preference.

The behavioral analyses of *npff ^-/-^* male and *gnrh3^-/-^* male demonstrated that NPFF and GnRH3 facilitate sexual behavior of male, which plays more active roles than female in medaka. On the other hand, both male and female of *gnrh3^-/-^*;*npff ^-/-^* showed normal sexual behavior to the WT partners of the opposite sex (Figure 3). In the previous behavioral studies of TN-GnRH neurons, male animals, whose TN-GnRH neurons were surgically ablated, showed a decrease in the motivation of sexual behavior (Yamamoto et al., 1997). This discrepancy may be caused by a possibility that surgical ablation damaged the other neurons than TN-GnRH neurons or that glutamate, which has been reported to coexist in these neurons (Akazome et al., 2011), have played a role in the control of motivation. In addition, since each single knockout (*npff ^-/-^* and *gnrh3^-/-^*) showed abnormal sexual behaviors, the possibility of developmental compensation in *gnrh3^-/-^*;*npff ^-/-^* is extremely low. We therefore propose a novel mechanism in which balanced release of NPFF and GnRH3 in the TN-GnRH neurons plays a key role in the control of male sexual behaviors, which is discussed below in detail (see *Balanced release of NPFF and GnRH3 in the TN-GnRH neurons may facilitate the motivation for male sexual behavior*).

### Anatomical distribution of NPFF and GnRH receptors suggests that NPFF and GnRH3 induce neuromodulation in the sexual behavior-controlling areas and the sensory processing areas

Next, based on the anatomical studies, we discuss the modulatory mechanism of NPFF and GnRH3. The present study and previous studies reported that TN-GnRH neurons express NPFF in addition to GnRH3 (Figre 1A, Wirsig-Wiechmann and Oka, 2002, Oehlmann et al., 2002, Saito et al., 2010). In teleosts, anatomical analysis using dwarf gourami demonstrated that a single TN-GnRH neuron projects their axons widely in the brain, from the olfactory bulb to the spinal cord (Oka and Matsushima, 1993). Since *npff ^-/-^* and *gnrh3^-/-^* showed the abnormal sexual behaviors in different manners (Figure 2 and 3), NPFF and GnRH3 are suggested to be released from TN-GnRH neurons to exert peptidergic neuromodulation to the specific receptor-expressing neurons, which is suggested to result in the increase in the motivation for sexual behaviors.

Our anatomical analyses demonstrated that NPFF receptors are expressed broadly in the brain (Figure 4A), especially in the brain regions controlling sexual behaviors (Vv, Vs, POA; Koyama et al., 1984, Satou et al., 1984). Previous study also reported that electrical stimulation of these brain regions induces male sexual behavior in hime salmon. Furthermore, also in mammals, NPFF receptors are expressed in the olfactory bulb and the lateral septum (Gouarderes et al., 2004), which are regions regulating sexual behaviors (Crews et al., 1996, Tobet and Baum, 1982). From these studies, it may be conserved among various vertebrates that NPFF receptors are expressed in the brain regions controlling sexual behaviors.

In addition, neurons expressing the immediate early gene *egr1* in the ventral part of POA (vPOA) were more abundantly observed in WT males than in *npff ^-/-^* males (Figure 6A and 6B). Around the time of removal of the transparent separation between the pairs, WT males are observed to be eagerly trying to mate with females, which means that the male’s motivation for sexual behavior is high. Actually, WT male spawned within 6 min after removing the transparent separation. Since the brain sampling was performed 25 – 28 min after removing the transparent separation, the *egr1* expression in the vPOA is suggested to be influenced by the motivation for the sexual behavior. Thus, the loss of *egr1* expression in the vPOA in *npff ^-/-^* may result in the lower motivation for male sexual behavior in these fish. On the other hand, in the analysis of co-expression of *egr1* and *gpr74-1*,which is the most predominant NPFF receptor subtype expressed in the vPOA (Figure 4D), the difference in the percentage of co-expressing neurons between WT and *npff ^-/-^* was not significant (Figure 6D). Thus, the majority of neurons showing the increase in *egr1*-expression in WT are suggested not to express NPFF receptors. Considering that there are neural circuitries among Vv, Vs, and POA (O’Connell and Hofmann, 2011) and local neural circuitries in POA (Zempo et al., 2018), NPFF shown to be expressed in TN-GnRH neurons in medaka (the present result) is suggested to modulate neurons in the vPOA via indirect pathway. Then, this neuromodulation may result in the increase in the motivation for male sexual behaviors. In the present study, we also showed that GnRH receptors are expressed mostly in the sensory processing areas (Figure 5A). Two subtypes of GnRH receptors, *gnrhr1* and *gnrhr3*, were expressed in the Dp, which is the olfactory and gustatory processing areas (Murakami et al., 1983, Yoshimoto et al., 1998). In addition, *gnrhr1*-expressing neurons were localized in the NI, which is the visual processing area (Ito et al., 1981). Moreover, the Dd and Dm, where *gnrhr3* was expressed, are somatosensory processing areas (Ito et al., 1986). Taken together, GnRH3 is suggested to modulate processing of multiple sensory modalities mainly via GnRHR1 and GnRHR3 in medaka. On the other hand, we found some *gnrhr3*-expressing neurons in the Vv and POA, where NPFF receptors were also expressed. It is suggested that GnRH3 modulates sexual behaviors via such brain regions.

Furthermore, in teleosts, TN-GnRH neurons project their axons to the inner granular layer in the retina, and neurons in this region project to the optic tectum, which suggests that TN-GnRH neurons integrate visual and olfactory information (Springer, 1983, Umino and Dowling, 1991, Li and Dowling, 2000, Li et al., 2017, Umatani and Oka, 2019). Considering these studies and our present results, NPFF and GnRH3 released from TN-GnRH neurons are suggested to integrate and modulate multiple sensory information and finally modulate the motivation for sexual behaviors.

### Balanced release of NPFF and GnRH3 in the TN-GnRH neurons may facilitate the motivation for male sexual behavior

We found that *gnrh3^-/-^* caused lower expression of *npff^-^* (Figure 7A). TN-GnRH neurons express GnRH receptors (Hajdu et al., 2007), and electrophysiological analyses demonstrated that GnRH modulates the electrical activities of TN-GnRH neurons in an auto/paracrine manner (Abe and Oka, 2000). Since GnRH-GnRHR signaling has been reported to regulate gene expressions (Naor, 2009), the present results may suggest GnRH3 regulation of NPFF expression in TN-GnRH neurons. To our knowledge, the present study is the first report suggesting expressional control of a peptide by another co-existing peptide, except for one paper reporting that a neuropeptide, cocaine- and amphetamine-regulated transcript, regulates the expression of co-existing neuropeptide, proopiomelanocortin (Lau et al., 2018). On the other hand, NPFF and GnRH3 are known to modulate neural activities by *in vitro* experiments (Karigo and Oka, 2013, Umatani and Oka, 2019), and their receptors were expressed in the sexual behavior-controlling areas (present results). Even though the expression of *npff* may be regulated by GnRH3, both NPFF and GnRH3 are suggested to be released from the terminal of TN-GnRH neurons and affect neurons in these brain areas. Based on the discussions above, we here hypothesize the following modulatory mechanism of male sexual behavior by TN-GnRH neurons (Figure 7B).

It should be noted that the function of NPFF and GnRH3 is neuromodulation of neural circuits controlling motivation for sexual behaviors and that the sexual behaviors *per se* can occur without these neuromodulations, because all the knockout medaka could eventually spawn in spite of some behavioral abnormalities. First, we made assumptions that GnRH3 facilitates NPFF expression and activates inhibitory neural circuits (blue). On the other hand, NPFF activates both inhibitory (blue) and excitatory neural circuits (pink) controlling male sexual behaviors. In WT, normal release of NPFF and GnRH3 causes balanced activities of excitatory and inhibitory neural circuits, which enables WT male to show normal sexual behaviors (Figure 7Ba). In *gnrh3^-/-^* medaka, loss of *gnrh3* expression and subsequently diminished expression of *npff* induce dominant activity of inhibitory neural circuits, which result in the low motivation for sexual behaviors (Figure 7Bb). In *npff ^-/-^*, loss of *npff* expression induces more dominant activity of inhibitory neural circuits and diminishes motivation for sexual behaviors (Figure 7Bc). On the other hand, in *gnrh3 ^-/-^*;*npff ^-/^*^-^ male, loss of both genes results in the balanced activity of inhibitory and excitatory neural circuits, which leads to the normal sexual behaviors (Figure 7Bd). In summary, our present study provides a novel mechanism by which balanced release of co-existing two peptides is important for the neuromodulatory function of TN-GnRH neurons, which may go a long way toward understanding the modulatory mechanism of motivation for sexual behaviors in response to external environments in general.

## Material and Methods

### Animal

We used male and female Japanese medaka (*Oryzias latipes*). They were kept at 27±1℃ under the following daylight condition: 14 h light (8:00–22:00) and 10 h dark. The fish were fed with artemia (Salt Creek Inc, Salt Lake, UT) or flake (Tetra, Yokohama, Japan) 2 – 5 times every day. We used sexually mature fish (> three months old), whose body weight was 0.25 ± 0.1g. All the experiments were conducted in accordance with the protocols approved by the animal care and use committee of Graduate School of Science, the University of Tokyo (Permission number, 15-3, 17-1, and 20-6).

### Generation of neuropeptide knockout medaka (*gnrh3 ^-/-^*, *npff ^-/-^*, *gnrh3 ^-/-^*;*npff ^-/-^*)

Knockout fish were generated by using transcription activator-like effector nuclease (TALEN) method. The cleavage site of each gene was designed by ZiFiT (supplied by ZINC FINGER CONSORTIUM; http://www.zincfingers.org/default2.htm). Each binding site of TALEN complex was the following:

gnrh3

Right - AGCCATCCATAGGACCA Left - GGCGTTGGTGGTTCAGG

npff

Right - GAGGCTGATGAAGAACA Left - GCTGCTGGGATTCCAGA

Transcription and injection of TALEN mRNA were performed as described in Takahashi *et al*., 2016 (Takahashi et al., 2016). We injected TALEN solution (25 ng/μl right- and left-TALEN mRNA, 7.5 ng/μl GFP mRNA, and 0.012% phenol red in phosphate buffered saline, PBS) into the cytoplasm of one or two cell-stage embryo. Homozygous transgenic offspring were obtained as described in Kayo et al., 2019 (Kayo et al., 2019). *gnrh3 ^-/-^*;*npff ^-/-^* medaka was generated by crossing *gnrh3 ^-/-^* medaka with *npff ^-/-^* medaka.

The genotype of these knockout medaka was examined by using PCR with LightCycler 96 (Roche Diagnostics, Basel, Switzerland). The genome DNA of each fish was extracted through proteinase K treatment (0.4 mg/ml proteinase K (Kanto Chemical Co., Inc., Tokyo, Japan) in TE (10 mM tris (hydroxymethyl) aminomethane-HCl 1mM, 1 mM Ethylenediaminetetraacetic acid, pH 8.0); 75℃ 20 min and 95℃ 10 min). Then, the genotype was identified by referring to the melting curve after PCR (95℃ 10 sec, 60℃ 10 sec, and 72℃ 10 sec; 45 cycles) and/or the direct sequencing after PCR (95℃ 20 sec, 70℃ 20 sec (−0.5℃/cycle), 72℃ 15 sec) × 20 cycles, (95℃ 20 sec, 60℃ 20 sec, 72℃ 15 sec) × 15 cycles). The sequences of primers are shown in Tables 1 and 2. The sequence of WT and each knockout were compared by using MAFFT (https://mafft.cbrc.jp/alignment/software/) or APE (https://jorgensen.biology.utah.edu/wayned/ ape/).

**Table 1.**
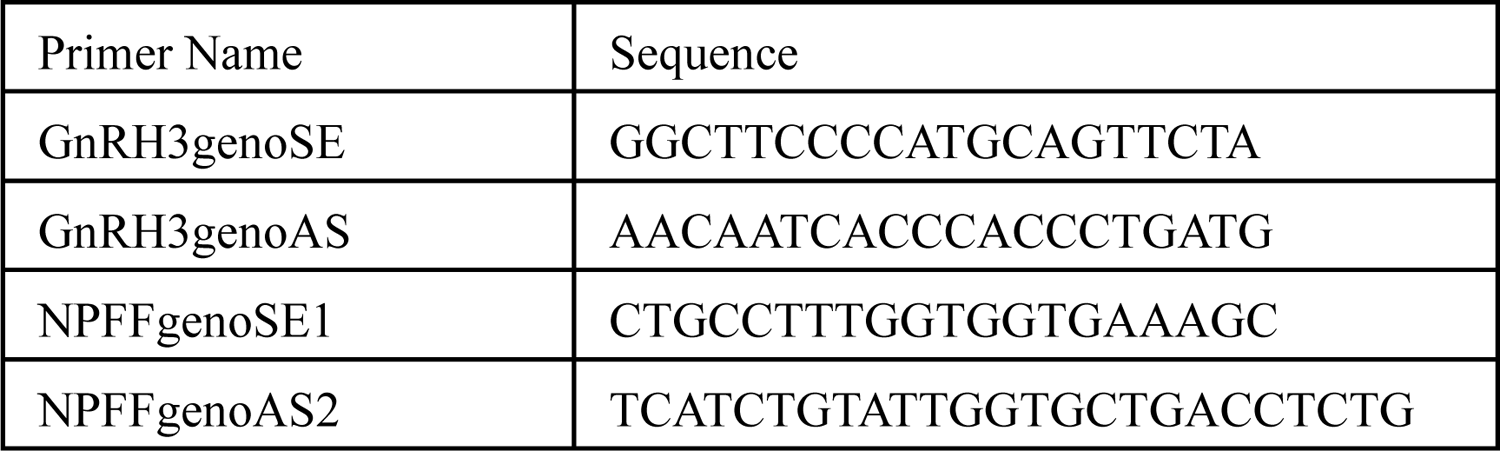
Primers for the melting curve analysis

**Table 2.**
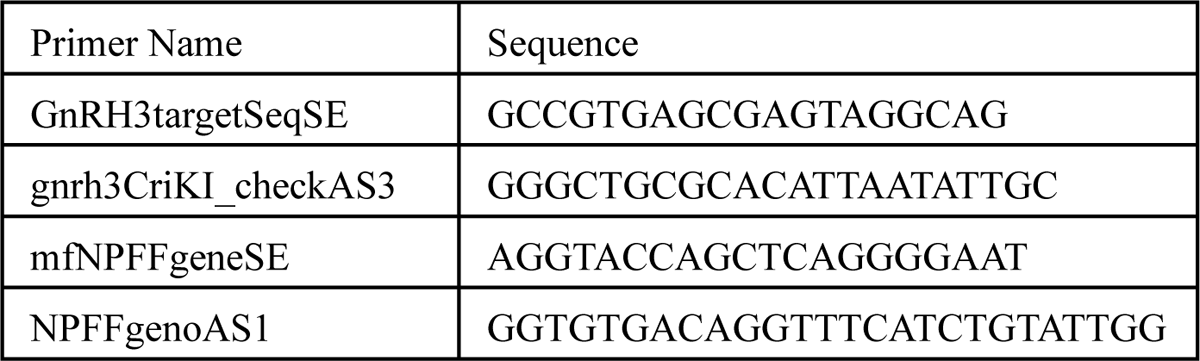
Primers for the sequence in order to examine genotype

### Immunohistochemistry (IHC)

To examine the loss of NPFF expression in *npff ^-/-^* medaka, we performed immunohistochemistry using FMRF-amide antibody (1:10000, FA1155, Enzo Life Sciences, Inc., Farmingdale, NY). In addition, we performed fluorescent IHC using GnRH3 specific antibody (LRH13, 1:10,000; (Park and Wakabayashi, 1986)) to examine the loss of GnRH3 expression in *gnrh3 ^-/-^* and *gnrh3 ^-/-^*;*npff ^-/-^* medaka. In each experiment, 25 µm frontal frozen sections were processed for the IHC procedure according to the standard protocol we documented previously (Karigo et al., 2014, Zempo et al., 2018). To analyze the cellular localizations and the axonal projections of TN-GnRH neurons, the 3,3’-Diaminobenzidine Tetrahydrochloride (DAB) -stained sections were counter-stained using the Nissl staining, after IHC using LRH13 antibody. For the nomenclature of the brain nuclei, we referred to Ishikawa *et al*., 1999 (Ishikawa et al., 1999).

### *In situ* hybridization (ISH)

Digoxigenin-labeled probes were made for each NPFF receptors (probe length; Gpr147: 1428 bases, Gpr74-1: 1323 bases, Gpr74-2: 1278 bases) and GnRH receptors (probe length; GnRHR1: 1234 bases, GnRHR2: 952 bases, GnRHR3: 1230 bases, GnRHR4: 1269bp). The brain sections were made following the methods described above (IHC). Hybridization, wash, and chromogenic reaction were performed as already reported in Zempo et al., 2013 (Zempo et al., 2013). After detection of signals by treatment with a chromogenic substrate, NBT/BCIP, slides were immersed in Neutral red solution (2 % Neutral red in distilled water: 2M phosphate buffer (pH4.8) = 1:1) for 3–8 min in order to visualize brain nuclei. Antisense-specific signals were identified by comparing with slides hybridized with sense probe.

For analyses of co-expression of *gnrh3*, *npff* and *gpr74-1*, a fluorescein-labelled probe for GnRH3 mRNA was prepared using fluorescein RNA labelling mix (Roche Diagnostics), and it was used in combination with the DIG-labelled RNA probes for *npff* or *gpr74-1* mRNA. After performing ISH procedures up to the blocking step described above, the sections were incubated with a horseradish peroxidase-conjugated anti-fluorescein antibody (diluted 1:500 with DIG-1, PerkinElmer, Foster City, CA) for 1 h and washed twice with DIG-1 for 5 min each time. The sections were then incubated with Biotinyl Tyramide Amplification kits (TSA, TSA plus biotin kit, PerkinElmer), following the manufacture’s protocol. After washing and signal amplification with avidin biotin complex reagents (1:100 in DIG-1; Vector Laboratories), they were next washed twice with DIG-1 for 5 min each, incubated with Alexa Fluor 488-conjugated streptavidin (diluted 1:500 in DIG-1; Thermo Fisher Scientific) and alkaline phosphatase-conjugated anti-DIG antibody (diluted 1:1,000 in DIG-1, Roche Diagnostics) for 2 h and washed twice with DIG-1 for 5 min each. After the detection of the GnRH3 fluorescent signal, alkaline phosphatase activity was detected using an HNPP fluorescence detection kit (Roche Diagnostics) according to the manufacturer’s instructions. Incubation for this substrate was conducted until visible signals were detected. After detection, the reaction was stopped by washing in PBS solution, and the sections were cover-slipped with CC/Mount (Diagnostic BioSystems, Pleasanton, CA).

For the analyses of co-expression of *egr1*and *gpr74-1*, a fluorescein-labelled probe for *gpr74-1* mRNA was prepared using a fluorescein RNA labelling mix, which was used in combination with the DIG-labelled RNA probes (1308 bases) for *egr1* mRNA. We prepared fish pairs according to the behavioural experiment procedure for *npff ^-/-^* (see below). Then, we separated the pairs of WT or *npff ^-/-^* in the evening before the experimental day and performed perfusion-fixation 25 – 28 min after removing the transparent separation in the next morning. After performing ISH procedures up to the blocking step described above, the sections were incubated with anti-DIG HRP antibody (Roche), and the signals for *egr1* were detected with TSA Cy3 kits (PerkinElmer). To block TSA reactions due to the activity of residual HRP carried over in the anti-DIG antibody, we incubated the sections in 3% H_2_O_2_ for 40 min. After washing, we detected signals for *gpr74-1* following the protocol for fluorescein-labelled *gnrh3* mRNA probe described above, with minor modification. After incubation with HRP-conjugated anti-fluorescein antibody, we performed TSA reaction (TSA plus biotin, Akoya Biosciences, Marlborough, MA) and ABC reaction (Vector laboratories) and detected signals for *gpr74-1* with Alexa Fluor 488-conjugated streptavidin. The cell nuclei were labelled by DAPI (Dojindo laboratories, Kumamoto, Japan). After washing in PBS solution, the sections were cover-slipped with FluoroMount/Plus (Diagnostic BioSystems).

We counted each labelled cells of three sections (thickness: 25 μm), which contain *gpr74-1*-expressing neurons in the vPOA, per one fish.

### Photomicroscopy

The photomicrographs of the fluorescence-labeled sections were taken with laser confocal microscopy (Zeiss LSM710; Carl Zeiss, Oberkochen, Germany). The photomicrographs of DAB-visualized or NBT/BCIP-visualized sections were taken by using a digital camera (DFC310FX; Leica Microsystems, Wetzlar, Germany) attached to a DM 5000 B microscope (Leica Microsystems).

### Behavioral analysis

For video recording of sexual behavior, we used a water tank with white sides and transparent bottom (15 cm × 15 cm × 12 cm). The test tank contained about 1 L water at 28℃. A single board computer Raspberry pi (Raspberry pi ZERO W; Raspberry Pi Foundation, Cambridge, UK) with a Raspberry Pi Camera V2 (Raspberry Pi Foundation) was placed under the tank. For behavioral analysis of phenotypes of *gnrh3 ^-/-^*, *npff ^-/-^*, and *gnrh3 ^-/-^*;*npff ^-/-^*, we paired one of these knockout medaka, either male or female, with WT of the opposite sex medaka. We kept the pairs in the same tank for 4–6 days before the behavioral recordings. Their behaviors were recorded one or two days after the pairs were moved to the test tank. We used WT pairs for control, which spawned eggs every day for successive three days. Each pair was separated by a transparent sheet with some slits during the night before the behavioral recordings. The behaviors were recorded from 9:50 AM and analyzed for 25 mins after removing the transparent separation at 10:00 AM.

We analyzed medaka behaviors by using the method described by Tomihara et al. (Tomihara et al., 2021). All the analyses were performed without knowing the genotype of the pair (blind test).

The parameters for behavioral analyses were as follows:

– latency to the first following (min); If a male did not follow a female at all in 25 min, the latency of that pair was defined as 25 min.
– latency to the first courtship (min); If a male did not show courtship at all in 25 min, the latency of that pair was defined as 25 min.
– latency to the first clasping (min); If a male did not show clasping at all in 25 min, the latency of that pair was defined as 25 min.
– latency to the spawning (min); If a pair did not spawn in 25 min, the latency of that pair was defined as 25 min.
– frequency of courtship before spawning (times/min); the frequency was calculated by dividing the number of courtships by the latency to spawning.
– percentage of following with successful courtship (%); This analysis excluded a pair, whose male did not show following at all.

### Cloning of medaka *gpr147* and *gpr74* (Npff receptors)

The cDNAs encoding medaka *gpr147* and *gpr74* were isolated, and their sequences were determined as previously described (Kanda et al., 2008). Briefly, total RNA of medaka brain was extracted from medaka using ISOGEN (Nippon Gene Co., Ltd., Tokyo, Japan), and polyadenylated RNA was enriched using Oligotex-dT30 Super mRNA Purification kits (Takara Bio Inc.). The cDNAs that were used for RACE were synthesised from 1 µg of polyA+ RNA using a SMART RACE cDNA Amplification Kit (Clontech Laboratories, Inc., Palo Alto, CA) according to the manufacturer’s instructions. Blasting the known NPFF receptor (GPR147 and GPR74) sequences against the medaka genome sequence database (Ensembl, http://www.ensembl.org/Multi/blastview), three candidate partial sequences (one for *gpr147* and two for *gpr74*) were obtained. In order to determine the sequences of both the 5′ and 3′ ends of medaka *gpr147* and *gpr74*, gene-specific oligonucleotide primers were designed and subjected to RACE. The sequences of the primers used are listed in Table 3. Polymerase chain reactions (PCR) were conducted in 10 µL containing 10× ExTaq buffer, dNTP, ExTaq polymerase (Takara Bio Inc.), 1 µL of template cDNA, 1× Universal Primer Mix (first PCR, Clontech Laboratories, Inc.) or 1× Nested Universal Primer (secondary PCR, Clontech Laboratories, Inc.) and gene-specific primers. The reaction conditions were as follows: 94 °C for 5 min; 5 cycles of 94 °C for 30 s and 72 °C for 2 min; 5 cycles of 94 °C for 30 s, 70 °C for 30 s, and 72 °C for 2 min; and 17 cycles of 94 °C for 30 s, 64 °C for 30 s, and 72 °C for 2 min. 200 nl of the first PCR solution was used for the next nested PCR. The reaction conditions for the nested PCR were as follows: 94 °C for 5 min; 5 cycles of 94 °C for 30 s and 72 °C for 2 min; 5 cycles of 94 °C for 30 s, 70 °C for 30 s, and 72 °C for 2 min; and 25 cycles of 94 °C for 30 s, 64 °C for 30 s, and 72 °C for 2 min. The amplified PCR products were purified on 1.5% agarose gels and cloned into the pGEM-T vector (Promega Corporation, Madison, WI) for sequence analysis. The nucleotide sequences were determined by analyzing more than three clones obtained from independent amplification to avoid PCR error. Transmembrane domains were predicted using the TMHMM server version 2.0.

**Table 3.**
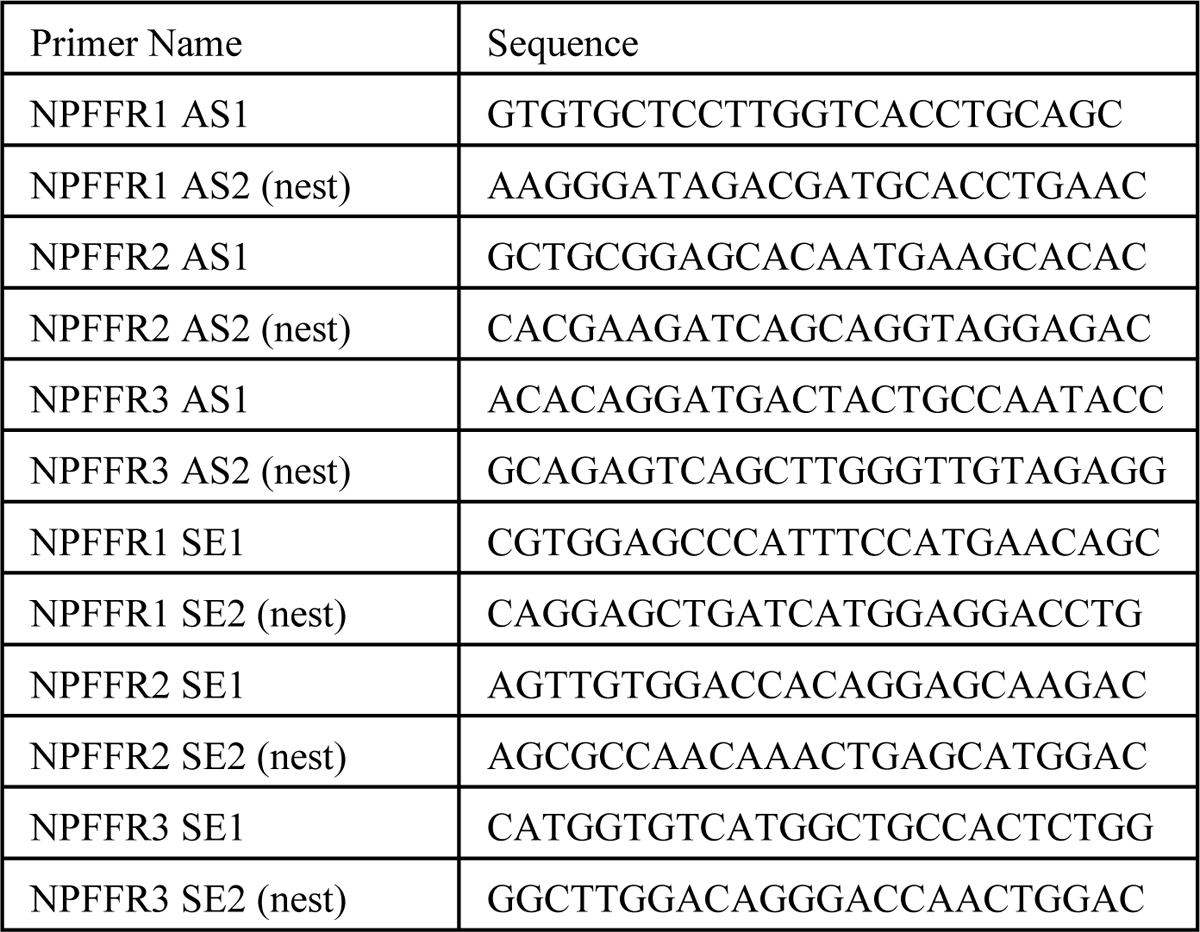
Primers for 5’ RACE and 3’ RACE (NPFFR1: Gpr147, NPFFR2: Gpr74-1, NPFFR3: Gpr74-2)

### Phylogenetic analysis

Multiple amino-acid sequence alignments were generated with Clustal X (2.1) (Thompson et al., 1997). The phylogenetic relationships among sequences were analysed with MEGA6 using the neighbour-joining method with 1000 bootstrap replicates (Tamura et al., 2007). The full-length amino acid sequences of GPR147 (NPFFR1) and GPR74 (NPFFR2) were retrieved from GenBank. GenBank accession numbers of used sequences are as follows: human NPFFR1 (AAK94199); cattle NPFFR1 (XP_612190); opossum GPR147 (XP_001374817); dog GPR147 (XP_851935); horse NPFFR1(XP_001917713); rat NPFFR1 (NP_071627); mouse RFRPR (XP_137119); pig NPFFR1 (XP_001925251); chicken Rfrpr (BAE17050); quail Gnihr (BAD86818); *Takifugu* Rfrpr (NP_001092117); zebrafish Gnihr1 (GU290219), Gnihr2 (GU290220), Gnihr3 (GU290221); human GPR74 (AAK58513); cattle NPFF2 (XP_609034); rat NPFF2 (NP_076470); mouse GPR74 (AAK58514); opossum NPFF2 (XP_001374952); chicken Npff2 (NP_001029997); zebrafish Npff2-1 (XP_693180); zebrafish Gpr74 (XP_690069); *Takifugu* Npff2-1 (NP_001092118); *Takifugu* Npff2-2 (NP_001092119).

### Luciferase assays

The cDNAs containing full-length open reading frames of Gpr147, Gpr74-1, Gpr74-2, and GnRHR4 were subcloned into the expression vector pcDNA3.1+ (Thermo fisher scientific). HEK293T cells were grown at 37 °C in Dulbecco’s modified Eagle’s medium supplemented with 5% foetal bovine serum. When cells reached 70-80% confluency in 10 cm dish, the medium was replaced with Opti-MEM (Thermo fisher scientific), and the plasmid DNAs (5.5 μg/dish) were transfected into monolayer culture cells with pGL4.29 containing CRE-luciferase reporter gene or pGL4.33 containing SRE-luciferase reporter gene (9.0 μg/dish, Promega) and pGL4.74 containing the Renilla luciferase reporter gene (0.5 μg/dish, Promega), as an internal control, using Polyethyleneimin “Max” (Polyscience, Inc., Warrington, PA) according to the manufacturer’s instructions. Cells were maintained in transfection mixture for 24 h. They were then moved to 24-well plates, treated with NPFF at various concentrations ranging 0 to 10^-5^M with or without 5 µM forskolin for 6 h before being harvested and analysed. Luminescence was measured with the Dual-Glo Luciferase Assay System (Promega) in a Lumat LB 9507 luminometer (EG & G, Berthold, Germany). Transfection experiments were performed in triplicate and repeated at least three times.

### Quantitative real-time PCR

We dissected a whole brain from WT, *gnrh3 ^-/-^*, or *npff ^-/-^* male. Total RNA of brain was extracted by using FastGene^TM^ RNA basic kit according to the manufacturer’s instructions. We performed reverse-transcription of 500 ng total RNA with FastGene^TM^ cDNA Synthesis 5× ReadyMix OdT (NIPPON Genetics Co., Ltd) according to the manufacturer’s instructions. For real time PCR, 1μl of cDNA diluted with 10-fold MQ was mixed with KAPA SYBR Fast qPCR kit and amplified with Lightcycler 96 (Roche; 95℃ 5 min, (95℃ 10 sec, 60℃ 10 sec, 72℃ 10 sec) × 45 cycles).

The data were normalized to a housekeeping gene, ribosomal protein s13.

### Data analysis

Sample sizes, as indicated in figure legends, were determined based on the observed variability across measurements. All experiments have been replicated in multiple independent trials (at least twice). All data were included in figures. For statistical analysis, we used Kyplot 5.0 software (Kyence, Tokyo, Japan) or R software (version 4.0.3, https://www.r-project.org/) with R studio (https://rstudio.com/products/rstudio/). Steel’s multiple comparison test (compared with WT pairs or WT) was used for multiple comparison of the latency of sexual behavior or the expression level. For the comparison of the percentage of pairs in which sexual behavior occurred, we used ‘survival’ package in R (https://cran.r-project.org/web/packages/survival/index.html) and Log- rank test with Bonferroni’s correction. For the comparison of the percentage of neurons between WT and *npff ^-/-^*, we used Mann-Whitney U test. The number of cells was shown as mean ± SD. In all statistical analysis, significance levels were described as follows; *: p < 0.05, **: p < 0.01, ***: p < 0.001.

## Data availability

The sequences of NPFF receptors were deposited in GenBank under the accession number AB697148 for GPR147, AB697149 for GPR74-1, and AB697150 for GPR74-2. All data are included in the manuscript and supporting source data files. Source data files have been provided for Figure 2, Figure 3, Figure4—figure supplement2, Figure5—figure supplement1, Figure 6, and Figure 7.

## Acknowledgements

We thank Dr. Nakajo (Osaka Medical and Pharmaceutical University) for helpful discussion and advice. We also thank Mr. Tomihara (the University of Tokyo) for his help with the behavioral analysis system. In addition, we thank Ms. Ruka Kojima for her construction of TALEN for knockout of *gnrh3* and Ms. Kiyoko Kataoka for her advice on *in situ* hybridization and her help in performing the double ISH. We also thank Ms. Hisako Kohno, Ms. Miho Kyokuwa, Ms. Fumika Muguruma, Ms. Fujiko Masui, Ms. Natsuko Ito, and Ms. Hiroko Tsukamoto for animal care. This work was supported by Grants-in-Aid from Japan Society for the Promotion of Science (JSPS) Grant (Grant number 17K15157 and 20H03071 to C. U., 24570067 to Y. A., 18H04881 and 18K19323 to S. K., and 26221104 to Y. O.), grant for Basic Science Research Projects from Sumitomo Foundation to S. K., and Research Grants in the Natural Sciences from the Mitsubishi Foundation to S. K.

## References

1. Abe, H. & Oka, Y. 2000. Modulation of pacemaker activity by salmon gonadotropin-releasing hormone (sGnRH) in terminal nerve (TN)-GnRH neurons. J Neurophysiol, 83, 3196–3200.

2. Akazome, Y., Kanda, S. & Oka, Y. 2011. Expression of vesicular glutamate transporter-2.1 in medaka terminal nerve gonadotrophin-releasing hormone neurones. J Neuroendocrinol, 23, 570–6.

3. Argiolas, A. & Melis, M. R. 2013. Neuropeptides and central control of sexual behaviour from the past to the present: a review. Prog Neurobiol, 108, 80–107.

4. Bonini, J. A., Jones, K. A., Adham, N., Forray, C., Artymyshyn, R., Durkin, M. M., Smith, K. E., Tamm, J. A., Boteju, L. W., Lakhlani, P. P., Raddatz, R., Yao, W. J., Ogozalek, K. L., Boyle, N., Kouranova, E. V., Quan, Y., Vaysse, P. J., Wetzel, J. M., Branchek, T. A., Gerald, C. & Borowsky, B. 2000. Identification and characterization of two G protein-coupled receptors for neuropeptide FF. J Biol Chem, 275, 39324–31.

5. Burmeister, S. S. & Fernald, R. D. 2005. Evolutionary conservation of the egr-1 immediate-early gene response in a teleost. J Comp Neurol, 481, 220–32.

6. Crews, D., Godwin, J., Hartman, V., Grammer, M., Prediger, E. A. & Sheppherd, R. 1996. Intrahypothalamic implantation of progesterone in castrated male whiptail lizards (Cnemidophorus inornatus) elicits courtship and copulatory behavior and affects androgen receptor- and progesterone receptor-mRNA expression in the brain. J Neurosci, 16, 7347–52.

7. Dennison, E., Bain, P. A., Bartke, A. & Meliska, C. J. 1996. Systemically administered gonadotrophin-releasing hormone enhances copulatory behaviour in castrated, testosterone-treated hyperprolactinaemic male rats. Int J Androl, 19, 253–9.

8. Dorsa, D. M. & Smith, E. R. 1980. Facilitation of mounting behavior in male rats by intracranial injections of luteinizing hormone-releasing hormone. Regul Pept, 1, 147–55.

9. Dorsa, D. M., Smith, E. R. & Davidson, J. M. 1981. Endocrine and behavioral effects of continuous exposure of male rats to a potent luteinizing hormone-releasing hormone (LHRH) agonist: evidence for central nervous system actions of LHRH. Endocrinology, 109, 729–35.

10. Gehlert, D. R., Thompson, L. K., Hemrick-Luecke, S. K. & Shaw, J. 2008. Monoaminergic compensation in the neuropeptide Y deficient mouse brain. Neuropeptides, 42, 367–75.

11. Gouarderes, C., Puget, A. & Zajac, J. M. 2004. Detailed distribution of neuropeptide FF receptors (NPFF1 and NPFF2) in the rat, mouse, octodon, rabbit, guinea pig, and marmoset monkey brains: a comparative autoradiographic study. Synapse, 51, 249–69.

12. Hajdu, P., Ikemoto, T., Akazome, Y., Park, M. K. & Oka, Y. 2007. Terminal nerve gonadotrophin-releasing hormone (GnRH) neurones express multiple GnRH receptors in a teleost, the dwarf gourami (Colisa lalia). J Neuroendocrinol, 19, 475–9.

13. Hiraki-Kajiyama, T., Yamashita, J., Yokoyama, K., Kikuchi, Y., Nakajo, M., Miyazoe, D., Nishiike, Y., Ishikawa, K., Hosono, K., Kawabata-Sakata, Y., Ansai, S., Kinoshita, M., Nagahama, Y. & Okubo, K. 2019. Neuropeptide B mediates female sexual receptivity in medaka fish, acting in a female-specific but reversible manner. Elife, 8.

14. Ishikawa, Y., Yoshimoto, M. & Ito, H. 1999. A brain atlas of a wild-type inbred strain of the medaka, Oryzias latipes. Fish Biology Journal Medaka, 10, 1–26.

15. Ito, H., Murakami, T., Fukuoka, T. & Kishida, R. 1986. Thalamic fiber connections in a teleost (Sebastiscus marmoratus): visual somatosensory, octavolateral, and cerebellar relay region to the telencephalon. J Comp Neurol, 250, 215–27.

16. Ito, H., Tanaka, H., Sakamoto, N. & Morita, Y. 1981. Isthmic afferent neurons identified by the retrograde HRP method in a teleost, Navodon modestus. Brain Res, 207, 163–9.

17. Kanda, S., Akazome, Y., Matsunaga, T., Yamamoto, N., Yamada, S., Tsukamura, H., Maeda, K. & Oka, Y. 2008. Identification of KiSS-1 product kisspeptin and steroid-sensitive sexually dimorphic kisspeptin neurons in medaka (oryzias latipes). Endocrinology, 149, 2467–76.

18. Karigo, T., Aikawa, M., Kondo, C., Abe, H., Kanda, S. & Oka, Y. 2014. Whole brain-pituitary in vitro preparation of the transgenic medaka (Oryzias latipes) as a tool for analyzing the differential regulatory mechanisms of LH and FSH release. Endocrinology, 155, 536–547.

19. Karigo, T. & Oka, Y. 2013. Neurobiological study of fish brains gives insights into the nature of gonadotropin-releasing hormone 1-3 neurons. Front Endocrinol (Lausanne), 4, 177.

20. Kawai, T., Abe, H., Akazome, Y. & Oka, Y. 2010. Neuromodulatory effect of GnRH on the synaptic transmission of the olfactory bulbar neural circuit in goldfish, *Carassius auratus*. J Neurophysiol, 104, 3540–50.

21. Kayo, D., Zempo, B., Tomihara, S., Oka, Y. & Kanda, S. 2019. Gene knockout analysis reveals essentiality of estrogen receptor beta1 (Esr2a) for female reproduction in medaka. Sci Rep, 9, 8868.

22. Kim, D. K., Cho, E. B., Moon, M. J., Park, S., Hwang, J. I., Kah, O., Sower, S. A., Vaudry, H. & Seong, J. Y. 2011. Revisiting the evolution of gonadotropin-releasing hormones and their receptors in vertebrates: secrets hidden in genomes. Gen Comp Endocrinol, 170, 68–78.

23. Kinoshita, M., Kobayashi, S., Urano, A. & Ito, E. 2007. Neuromodulatory effects of gonadotropin-releasing hormone on retinotectal synaptic transmission in the optic tectum of rainbow trout. Eur J Neurosci, 25, 480–4.

24. Koyama, Y., Satou, M., Oka, Y. & Ueda, K. 1984. Involvement of the telencephalic hemispheres and the preoptic area in sexual behavior of the male goldfish, Carassius auratus: a brain-lesion study. Behav Neural Biol, 40, 70–86.

25. Lau, J., Farzi, A., Qi, Y., Heilbronn, R., Mietzsch, M., Shi, Y. C. & Herzog, H. 2018. CART neurons in the arcuate nucleus and lateral hypothalamic area exert differential controls on energy homeostasis. Mol Metab, 7, 102–118.

26. Li, L. & Dowling, J. E. 2000. Disruption of the olfactoretinal centrifugal pathway may relate to the visual system defect in *night blindness b* mutant zebrafish. J Neurosci, 20, 1883–92.

27. Li, L., Wojtowicz, J. L., Malin, J. H., Huang, T., Lee, E. B. & Chen, Z. 2017. GnRH-mediated olfactory and visual inputs promote mating-like behaviors in male zebrafish. PLoS One, 12, e0174143.

28. Murakami, T., Morita, Y. & Ito, H. 1983. Extrinsic and intrinsic fiber connections of the telencephalon in a teleost, Sebastiscus marmoratus. J Comp Neurol, 216, 115–31.

29. Myers, B. M. & Baum, M. J. 1980. Facilitation of copulatory performance in male rats by naloxone: effects of hypophysectomy, 17 alpha-estradiol, and luteinizing hormone releasing hormone. Pharmacol Biochem Behav, 12, 365-70.

30. Naor, Z. 2009. Signaling by G-protein-coupled receptor (GPCR): studies on the GnRH receptor. Front Neuroendocrinol, 30, 10–29.

31. O’connell, L. A. & Hofmann, H. A. 2011. The vertebrate mesolimbic reward system and social behavior network: a comparative synthesis. J Comp Neurol, 519, 3599–639.

32. Oehlmann, V. D., Korte, H., Sterner, C. & Korsching, S. I. 2002. A neuropeptide FF-related gene is expressed selectively in neurons of the terminal nerve in Danio rerio. Mechanisms of Development, 117, 357–361.

33. Oka, Y. & Matsushima, T. 1993. Gonadotropin-releasing hormone (GnRH)-immunoreactive terminal nerve cells have intrinsic rhythmicity and project widely in the brain. J Neurosci, 13, 2161–2176.

34. Okubo, K., Ishii, S., Ishida, J., Mitani, H., Naruse, K., Kondo, M., Shima, A., Tanaka, M., Asakawa, S., Shimizu, N. & Aida, K. 2003. A novel third gonadotropin-releasing hormone receptor in the medaka Oryzias latipes: evolutionary and functional implications. Gene, 314, 121–131.

35. Okubo, K. & Nagahama, Y. 2008. Structural and functional evolution of gonadotropin-releasing hormone in vertebrates. Acta Physiol (Oxf), 193, 3–15.

36. Okuyama, T., Yokoi, S., Abe, H., Isoe, Y., Suehiro, Y., Imada, H., Tanaka, M., Kawasaki, T., Yuba, S., Taniguchi, Y., Kamei, Y., Okubo, K., Shimada, A., Naruse, K., Takeda, H., Oka, Y., Kubo, T. & Takeuchi, H. 2014. A neural mechanism underlying mating preferences for familiar individuals in medaka fish. Science, 343, 91–4.

37. Ono, Y. & Uematsu, T. 1957. Mating ethogram in *oryzias latipes*. Jour Fac Sci Hokkaido Univ Ser VI Zool, 13, 197–202.

38. Park, M. K. & Wakabayashi, K. 1986. Preparation of a monoclonal antibody to common amino acid sequence of LHRH and its application. Endocrinol Jpn, 33, 257–72.

39. Saito, T. H., Nakane, R., Akazome, Y., Abe, H. & Oka, Y. 2010. Electrophysiological analysis of the inhibitory effects of FMRFamide-like peptides on the pacemaker activity of gonadotropin-releasing hormone neurons. J Neurophysiol, 104, 3518–3529.

40. Satou, M., Oka, Y., Kusunoki, M., Matsushima, T., Kato, M., Fujita, I. & Ueda, K. 1984. Telencephalic and preoptic areas integrate sexual behavior in hime salmon (landlocked red salmon, Oncorhynchus nerka): results of electrical brain stimulation experiments. Physiol Behav, 33, 441–7.

41. Springer, A. 1983. Centrifugal innervation of goldfish retina from ganglion-cells of the nervus terminalis. Journal of Comparative Neurology, 214, 404–415.

42. Takahashi, A., Kanda, S., Abe, T. & Oka, Y. 2016. Evolution of the Hypothalamic- Pituitary-Gonadal Axis Regulation in Vertebrates Revealed by Knockout Medaka. Endocrinology, 157, 3994–4002.

43. Tamura, K., Dudley, J., Nei, M. & Kumar, S. 2007. MEGA4: Molecular Evolutionary Genetics Analysis (MEGA) software version 4.0. Mol Biol Evol, 24, 1596–9.

44. Thompson, J. D., Gibson, T. J., Plewniak, F., Jeanmougin, F. & Higgins, D. G. 1997. The CLUSTAL_X windows interface: flexible strategies for multiple sequence alignment aided by quality analysis tools. Nucleic Acids Res, 25, 4876–82.

45. Tobet, S. A. & Baum, M. J. 1982. Implantation of dihydrotestosterone propionate into the lateral septum inhibits sexual receptivity in estrogen-primed, ovariectomized rats. Neuroendocrinology, 34, 333–8.

46. Tomihara, S., Oka, Y. & Kanda, S. 2021. Establishment of open-source semi-automated behavioral analysis system and quantification of the difference of sexual motivation between laboratory and wild strains. Scientific Reports, 11, 10894.

47. Umatani, C., Misu, R., Oishi, S., Yamaguchi, K., Abe, H. & Oka, Y. 2015. GnRH suppresses excitability of visual processing neurons in the optic tectum. J Neurophysiol, 114, 2775–84.

48. Umatani, C. & Oka, Y. 2019. Multiple functions of non-hypophysiotropic gonadotropin releasing hormone neurons in vertebrates. Zoological Lett, 5, 23.

49. Umino, O. & Dowling, J. E. 1991. Dopamine release from interplexiform cells in the retina: effects of GnRH, FMRFamide, bicuculline, and enkephalin on horizontal cell activity. J Neurosci, 11, 3034–46.

50. Vilim, F. S., Aarnisalo, A. A., Nieminen, M. L., Lintunen, M., Karlstedt, K., Kontinen, V. K., Kalso, E., States, B., Panula, P. & Ziff, E. 1999. Gene for pain modulatory neuropeptide NPFF: induction in spinal cord by noxious stimuli. Mol Pharmacol, 55, 804–11.

51. Walter, R. O. & Hamilton, J. B. 1970. Head-up Movements as an Indicator of Sexual Unreceptivity in Female Medaka, *Oryzias latipes*. Animal Behaviour, 18, 125–127.

52. Wirsig-Wiechmann, C. R. & Oka, Y. 2002. The terminal nerve ganglion cells project to the olfactory mucosa in the dwarf gourami. Neurosci Res, 44, 337–41.

53. Wirsig, C. R. & Leonard, C. M. 1987. Terminal nerve damage impairs the mating behavior of the male hamster. Brain Res, 417, 293–303.

54. Yamamoto, N. & Ito, H. 2000. Afferent sources to the ganglion of the terminal nerve in teleosts. J Comp Neurol, 428, 355–75.

55. Yamamoto, N., Oka, Y. & Kawashima, S. 1997. Lesions of gonadotropin-releasing hormone-immunoreactive terminal nerve cells: effects on the reproductive behavior of male dwarf gouramis. Neuroendocrinology, 65, 403–12.

56. Yang, H. Y., Tao, T. & Iadarola, M. J. 2008. Modulatory role of neuropeptide FF system in nociception and opiate analgesia. Neuropeptides, 42, 1–18.

57. Yoshimoto, M., Albert, J. S., Sawai, N., Shimizu, M., Yamamoto, N. & Ito, H. 1998. Telencephalic ascending gustatory system in a cichlid fish, Oreochromis (Tilapia) niloticus. J Comp Neurol, 392, 209–26.

58. Zempo, B., Kanda, S., Okubo, K., Akazome, Y. & Oka, Y. 2013. Anatomical distribution of sex steroid hormone receptors in the brain of female medaka. J Comp Neurol, 521, 1760–80.

59. Zempo, B., Karigo, T., Kanda, S., Akazome, Y. & Oka, Y. 2018. Morphological Analysis of the Axonal Projections of EGFP-Labeled Esr1-Expressing Neurons in Transgenic Female Medaka. Endocrinology, 159, 1228–1241.

